# Comprehensive Metabolomic Analysis of Human Heart Tissue Enabled by Parallel Metabolite Extraction and High-Resolution Mass Spectrometry

**DOI:** 10.1101/2023.09.15.558013

**Authors:** Benjamin Wancewicz, Melissa R. Pergande, Yanlong Zhu, Zhan Gao, Zhuoxin Shi, Kylie Plouff, Ying Ge

**Affiliations:** Department of Cell and Regenerative Biology, University of Wisconsin-Madison, Madison, Wisconsin, 53705, USA; Human Proteomics Program, School of Medicine and Public Health, University of Wisconsin-Madison, Madison, Wisconsin, 53705, USA

**Keywords:** metabolomics, mass spectrometry, Fourier-transform ion cyclotron resonance, liquid chromatography mass spectrometry, heart metabolites

## Abstract

The heart contracts incessantly and requires a constant supply of energy, utilizing numerous metabolic substrates such as fatty acids, carbohydrates, lipids, and amino acids to supply its high energy demands. Therefore, a comprehensive analysis of various metabolites is urgently needed for understanding cardiac metabolism; however, complete metabolome analyses remain challenging due to the broad range of metabolite polarities which makes extraction and detection difficult. Herein, we implemented parallel metabolite extractions and high-resolution mass spectrometry (MS)-based methods to obtain a comprehensive analysis of the human heart metabolome. To capture the diverse range of metabolite polarities, we first performed six parallel liquid-liquid extractions (three monophasic, two biphasic, and one triphasic extractions) of healthy human donor heart tissue. Next, we utilized two complementary MS platforms for metabolite detection - direct-infusion ultrahigh-resolution Fourier-transform ion cyclotron resonance (DI-FTICR) and high-resolution liquid chromatography quadrupole time-of-flight tandem MS (LC-Q-TOF MS/MS). Using DI-FTICR MS, 9,521 metabolic features were detected where 7,699 were assigned a chemical formula and 1,756 were assigned an annotated by accurate mass assignment. Using LC-Q-TOF MS/MS, 21,428 metabolic features were detected where 626 metabolites were identified based on fragmentation matching against publicly available libraries. Collectively, 2276 heart metabolites were identified in this study which span a wide range of polarities including polar (benzenoids, alkaloids and derivatives and nucleosides) as well as non-polar (phosphatidylcholines, acylcarnitines, and fatty acids) compounds. The results of this study will provide critical knowledge regarding the selection of appropriate extraction and MS detection methods for the analysis of the diverse classes of human heart metabolites.

**Table of Contents Graphical Abstract:** 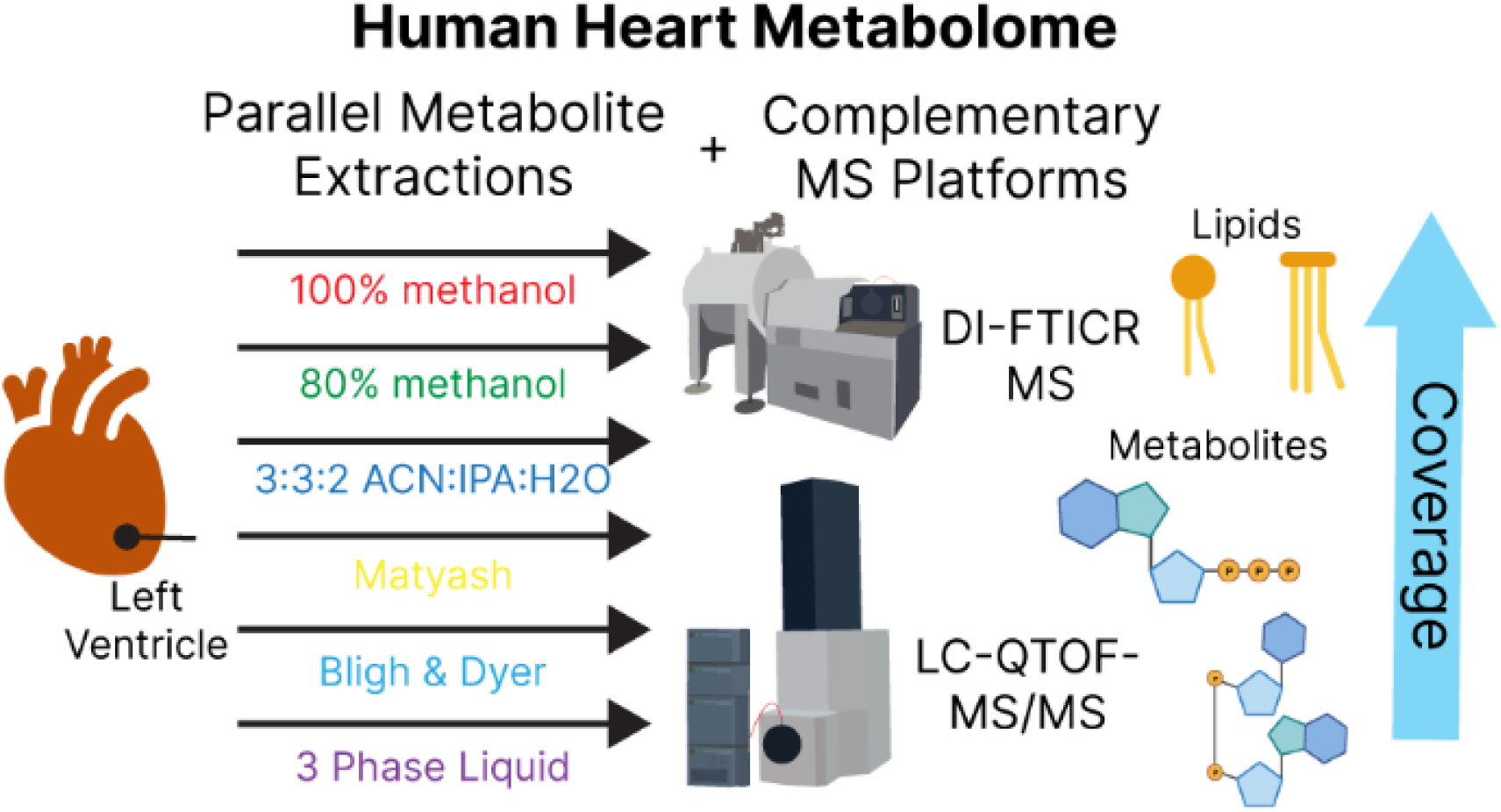

## Introduction

The heart contracts incessantly to supply oxygenated blood to the entire body and thus, requires a constant source of energy to fuel its high energy demands.^1^ The heart is a “metabolic omnivore” and utilizes a variety of fuel substrates, ranging from very polar (e.g., amino acid and nucleosides) to very non-polar (e.g., lipids).^2–6^ Therefore, a comprehensive metabolomic analysis is critical to understand cardiac metabolism; however, it remains challenging as the broad range of metabolite polarities makes extraction and detection difficult.^7–14^

First it is critical to effectively extract metabolites in a comprehensive and reproducible manner.^15, 16^ Generally, a monophasic extraction is the simplest, most reproducible way to enrich polar or non-polar compounds within a similar polarity range.^17^ In contrast, multiphasic extractions (e.g., biphasic or triphasic extractions) can enrich a broader range of polar and non-polar metabolites,^18^ thereby increasing metabolome coverage-one caveat being amphiphilic metabolite partitioning which can concomitantly introduce additional variability.^19^ In general, maximizing the metabolome coverage is essential to better understand cardiac metabolism.

Second, it is essential to efficiently detect and identify the extracted metabolites. There are multiple MS-based platforms that can be used for the analysis of metabolites, each having its own unique advantage.^20–22^ As previously demonstrated, direct infusion workflows can rapidly and reproducibly detect hundreds of metabolites when coupled with ultrahigh-resolution mass spectrometery.^23, 24^ On the other hand, there are many pathways which contain isomeric metabolites (e.g., isocitrate and citrate) which requires the implementation of additional separation strategies prior to MS detection. As a complementary technique, liquid chromatography (LC) separation coupled to high-resolution mass spectrometry allows for the separation of structural isomers while providing high-resolution detection.^22, 25^ Moreover, tandem spectra can be generated using this approach, allowing for fragmentation matching. The drawbacks of LC separation are longer analysis time and chromatographic bias of potential metabolites that can be detected.^22^ Therefore, utilization of both MS detection platforms would leverage the strengths of each technique and allow for a more comprehensive metabolomic analysis.

In this study, we combined parallel liquid-liquid metabolite extractions and complementary high-resolution MS detection methods to obtain a comprehensive analysis of the human heart metabolome. To capture the diverse range of metabolite polarities, we first performed six parallel liquid-liquid extractions of healthy human donor heart tissue - three monophasic, two biphasic, and one triphasic extractions. Next, analysis of extracted metabolites was performed using both direct infusion Fourier-transform ion cyclotron resonance (DI-FTICR) MS and liquid chromatography quadrupole time-of-flight tandem MS (LC-Q-TOF MS/MS). Collectively, we identified 2276 metabolites where each different extraction provides unique metabolite identifications. These included both previously reported and newly identified human heart metabolites. Additionally, pathway analysis of identified metabolites revealed increased coverage by using this comprehensive approach compared to any single extraction alone. Together, our findings underpin the importance of using parallel extractions and complementary MS detection methods in human heart metabolome studies and serve as a guide to assist in the selection of an appropriate extraction and MS method(s) to investigate specific metabolite classes.

## Experimental Section

### Chemicals and reagents

All reagents were purchased from Millipore Sigma (St Louis, MO, USA) and Fisher Scientific (Fair Lawn, NJ, USA) unless noted otherwise.

### Human Heart Tissue

All human subjects research was conducted in accordance with the standards set out by the World Medical Association (WMA) Declaration of Helsinki *Ethical Principles for Medical Research Involving Human Subjects* as approved by the University of Wisconsin (UW) Health Sciences Institutional Review Board (IRB) (UW 2013-1082).

### Metabolite extractions

Metabolite extraction was performed using 50mg of cryopulverized human heart tissue for all six types of extractions. For the single phase extractions, 1mL of ice-cold (−20°C) extraction solvent was added (100% methanol (MeOH), 80% MeOH, and 3:3:2 acetonitrile: isopropanol: water (ACN:IPA:H_2_O)). The tissue was homogenized using a Teflon pestle (1.5 mL microcentrifuge tube, flat tip, Thomas Scientific, Swedesboro, NJ, USA), vortexed, and placed on rotating nutator for 10 min. For the biphasic extractions, 225 µL MeOH was added for the Matyash extraction and 334 µL of MeOH for the Bligh & Dyer (B&D) extraction. Tissues were homogenized using a Teflon pestle, vortexed, and placed on a rotating nutator for 10 min. Next, 750 µL of methyl tert-butyl ether (MTBE) was added to the Matyash mixture and 300 µL of chloroform was added for B&D extraction mixture. To induce phase separation, 188 µL of H_2_O was added to the Matyash mixture and 267 µL of H_2_O to the B&D mixture. For the three-phase liquid extraction (3PLE), 250 µL of hexane was added to the tissue, homogenized using a Teflon pestle, vortexed, and placed on a rotating nutator for 10 min. Next, 250 µL methyl acetate, 225 µL acetonitrile, and 250 µL H_2_O were then added and vortexed. All samples were incubated for 10 min on a rotating nutator after stepwise addition of solvent. All samples were centrifuged for 2 min at 21,000 *g*, supernatant removed for each phase and dried *in vacuo*. Each of the different extractions were performed in triplicate.

### DI-FTICR MS analysis of metabolite extracts

FTICR MS detection of metabolites was performed using direct infusion by syringe pump into a Bruker solariX 12T FTICR mass spectrometer (Bruker Daltonics, Bremen, Germany) with the same mass spectrometry conditions as previously used.^23^ Full methods are available in the supplemental section. In brief, dried extracts were resuspended in 50:50 MeOH:H_2_O or 9:1 IPA:ACN for polar and nonpolar analysis, respectively. Samples were infused at 180 µL/hr for 300 scans into the FTICR MS which analyzed from *m/z* 50 to 1500. FTICR mass spectra were processed using ftmsProcessing software V2.3.0 (Bruker Daltonics, Bremen, Germany) to remove Gibbs and harmonic peaks. Bucket (mass) lists in positive and negative ion modes were generated using the T-ReX 2D workflow with a maximum mzDelta of 2 mDa, maximum charge state of 2, intensity threshold of 5E7, and the minimum number of features for the result was set to 2 of 3 replicates for a single phase. Positive and negative mode bucket lists were merged with a 1.0 ppm *m/z* tolerance. The merged bucket list was annotated with the SmartFormula function with 1.0 ppm as the narrow Δ*m/z* cutoff, 2.0 ppm as the wide Δ*m/z* cutoff, 20 as the narrow mSigma cutoff, and 50 as the wide mSigma cutoff. Putative metabolites were annotated using “hierarchical search”, which selects annotation from first library hit, against target lists containing previously detected metabolites with our platform, then outside libraries were considered-LipidBlast, internal MS/MS libraries, and Massbank of North America (MoNA) export for accurate mass matching with a 2.0 ppm mass error cutoff.

### LC-QTOF MS/MS analysis of metabolite extracts

Metabolite extracts were also analyzed via tandem mass spectrometry on an Impact II Q-TOF (Bruker Daltonics, Bremen, Germany) in both positive and negative ion modes. Detailed information on chromatographic conditions is provided in the SI. Given the different chemical properties of the metabolites in each extraction phase, the most suitable LC method(s) were paired with each extraction phase to maximize metabolome coverage. All analysis methods were 20 min and separation of metabolites was performed using a Waters nanoACQUITY system. Mass detection was performed from m*/z* 50 to 1300 at a spectral scan rate of 8Hz for MS1 with the top 5 precursors being selected for tandem (MS/MS) analysis employing a threshold of 500 counts. Ion source settings were as follows: end plate offset of 500 V, capillary voltage 4500 V, nebulizer at 0.5 bar and 4.0 L/min drying gas, and drying temperature of 200°C. Collision energies were 20, 25, and 30 eV for *m/z* 100, 500, and 1000. Raw data were analyzed using DataAnalysis 2022b. The settings for peak selection were peak area of 1000 for peak detection, minimum number of features in 2 of 3 replicates, minimum peak length of 8 spectra, and recursive peak length of 5 spectra. Primary ions were [M+H] ^+^, [M+Na] ^+^, [M+K] ^+^, [M+NH4] ^+^, [M-H] ^-^, and [M+HAc-H] ^-^, while [M+H-H_2_O] ^+^ was a common ion. Positive and negative mode peak lists were combined. Identifications were assigned from LipidBlast, internal MS/MS libraries, and Massbank of North America (MoNA) exports in parallel search with highest scoring match selected as annotation. Identifications had to have a MS/MS score of ≥400 and mass tolerance of 20 ppm.

### Data processing and analysis

Data output was manually reviewed, filtered, and chemical annotations added for identified metabolites. Metabolic features with a ratio of sample average: blank average < 10 were not reported. Also, metabolite features not present in 2 of 3 extraction replicates in at least one extraction were not reported. InChI key, HMDB ID, KEGG ID, and SMILES information was added for identified metabolites using The Chemical Translation Service (Fiehn laboratory, University of California Davis) or manually retrieved. InChI keys were used for batch ClassyFire analysis to get chemical class information.^26^. Pathway analysis was done using MetaboAnalyst 5.0.^27^ ChemRich plot was made using R script.^28^ Correlation analyses were generated using GraphPad Prism (v9.1). UpSet plot was made using UpSet R shiny web application.^29^

## Results and Discussion

### Parallel Extractions Maximizes Metabolome Coverage

In this study, we utilized a parallel extraction (three monophasic, two biphasic, and one triphasic extraction) approach to capture the broad range of metabolites in the human heart (**Figure 1**). For the monophasic extractions, we used 100% methanol (MeOH), 80% MeOH, and 3:3:2 acetonitrile:isopropanol:water (ACN:IPA:H_2_O). These extractions cover the majority of metabolite classes and are fast with the 3:3:2 ACN:IPA:H_2_O extraction covering the widest range, while the MeOH extraction is more nonpolar and the 80% MeOH being more polar.^25^ Due to their simplicity, sample throughput is higher, but lack in-depth extraction efficiency.^25^ Multiphasic approaches have proven useful for the selective extraction of lipids into one phase while partitioning more polar compounds in the aqueous phase.^30–32^ For biphasic extractions, we used the Bligh & Dyer (B&D)^30^ and Matyash^31^ extractions. Markedly, triphasic extractions (three phase liquid extraction, 3PLE) further separates the non-polar phase into polar and neutral lipid phases, retaining the polar metabolites in the aqueous phase.^32^ In general, multiphasic extractions enable a broader metabolome coverage compared to any monophasic extraction alone; one caveat of this approach is that some metabolites can partition into both phases, making metabolite detection and quantification potentially difficult. ACars are one example of this particular phenomenon; species with shorter fatty acyl chains mainly partition into the polar phase, while longer chains (>C10) species partition into the more non-polar phase.^19^ While separating lipids and polars metabolites into different phases requires additional analysis time, multiphasic extractions can be beneficial in situations where ion suppression is a concern, such as direct infusion MS analysis.

**Figure 1.**
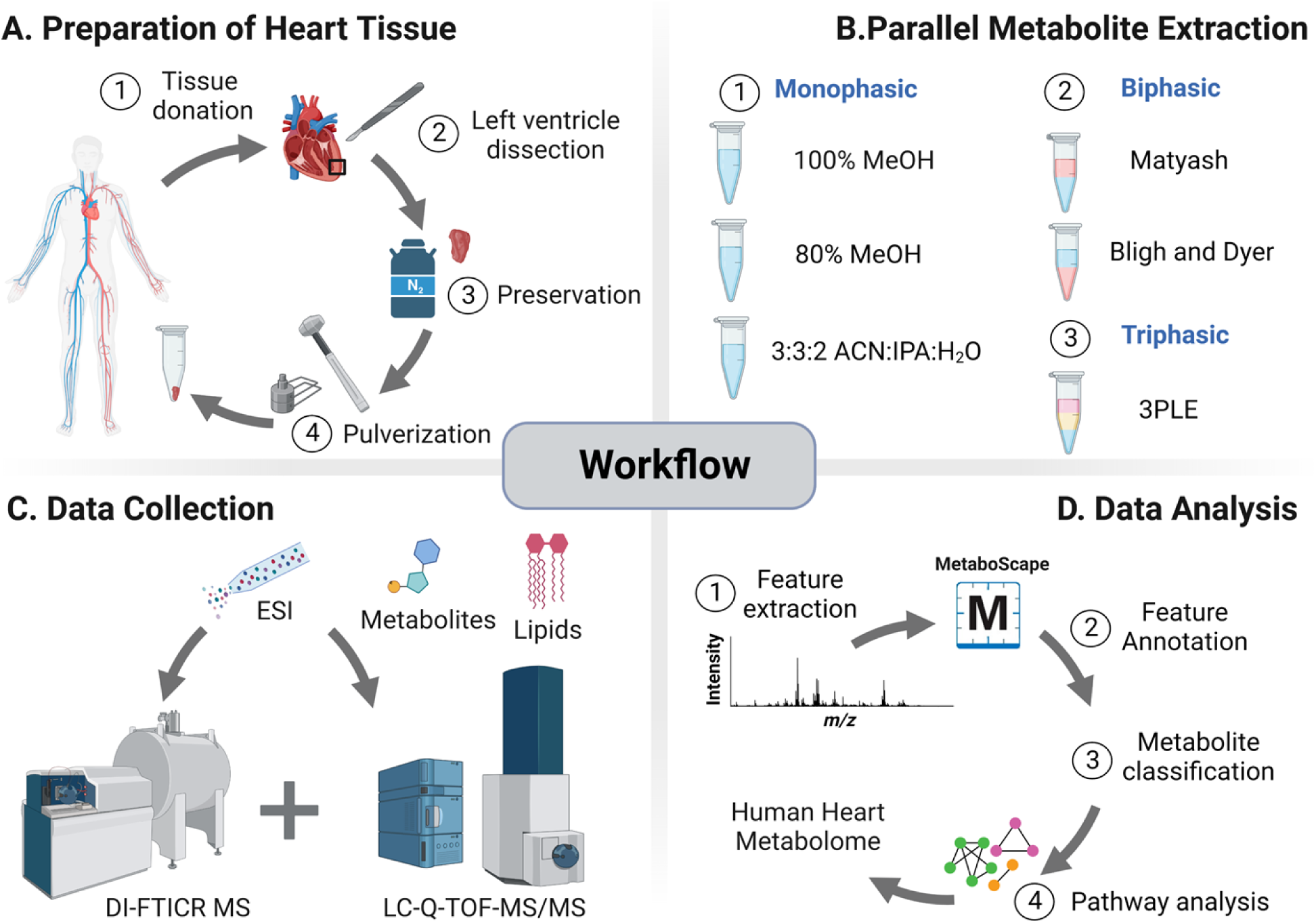
Overview of experimental workflow. (A) Left ventricle tissue from healthy donor tissue was dissected, snap frozen in liquid nitrogen, and cryopulverized. (B) Metabolites (polar metabolites and lipids) were extracted using multiple solvent mixtures including monophasic (100% MeOH, 80% MeOH and 3:3:2 ACN: IPA: H2O), biphasic (Matyash and Bligh and Dyer) and triphasic (3 phase liquid extraction, 3PLE) extraction methods; all extractions were done in triplicate. (C) Extracted metabolites were analyzed using ultrahigh-resolution mass spectrometry direct infusion Fourier-transform ion cyclotron resonance (DI-FTICR, Bruker SolariX 12T) and high-resolution liquid chromatography quadrupole time-of-flight tandem mass spectrometry (LC-Q-TOF MS/MS, Bruker Impact II) (D) Raw mass spectrometry data was analyzed where metabolic features were extracted and annotated. Further, metabolite classification and pathway analysis were performed to determine biological relevance. MeOH = methanol, ACN= acetonitrile, IPA= 2-propanol and H2O= water. Figure created with *BioRender.com*.

To compare the performance of the different extractions, we utilized DI-FTICR MS given its ultrahigh-resolution and robust reproducibility.^23^ Here, we evaluated metabolite enrichment reproducibility across all six extraction methods. First, we performed a visual inspection of MS spectra and observed it to be highly reproducible across three 100% MeOH extraction replicates in both positive and negative ion modes (Figure S1). Pearson correlation analysis of replicates was performed for all extraction phases: monophasic extractions (Figure S2), biphasic extractions (Figure S3), and triphasic extractions (Figure S4) as well as summarized by extraction method (Figure S5). For the monophasic extraction, 3:3:2 ACN: IPA: H2O performed the best with a Pearson correlation coefficient ranging from 0.9828 to 0.9972 across replicate comparisons. For the biphasic extractions, the Matyash extraction was observed to be slightly more reproducible than the B&D extraction with Pearson correlation coefficients ranging from 0.9664 to 0.9866 across replicate comparisons. Notably, the polar phase for B&D had better reproducibility compared to the non-polar phase, presumably contributing to the divergence between the two biphasic extractions (Figure S3). Interestingly, the lower phases had a lower Pearson correlation compared to the upper phases for both biphasic extractions (Figure S3). This additional variability indicates the potential difficulties with pipetting across multiple layers.

When examining the metabolome coverage, there was an increasing number of metabolic features observed with increasing number of extraction phases, wherein the triphasic extraction had the most metabolic features, SmartFormula (or chemical formula) annotations, and identified metabolites (**Figure 2A**). The number of SmartFormula annotations was compared separately among the monophasic, biphasic, and triphasic extractions to evaluate overlap. Here, 554, 488, and 840 SmartFormula annotations were uniquely detected in the 100% MeOH, 80% MeOH, and 3:3:2 ACN: IPA: H_2_0 extractions, respectively (**Figure 2B**). When comparing the overlap of the biphasic extractions, 848 and 414 SmartFormula annotations were detected only in the Matyash and B&D extractions, respectfully. Of these SmartFormula annotations 270, 343, 145, and 505 were observed to be detected only in the B&D polar phase, Matyash polar phase, B&D non-polar phase, and B&D polar phase, respectively (**Figure 2C**). When considering the triphasic extraction, 2361, 553, and 471 unique SmartFormula annotations were observed only in the polar, polar lipid, and non-polar lipid phases, respectively (**Figure 2D**). In the triphasic extraction, the majority of the unique SmartFormula annotations were observed in the polar phase, while the polar and non-polar lipid phases revealed a greater overlap. Comparing the overlap between the six extractions, the three largest intersection areas were: 1366 shared between all six extractions, 1063 annotations unique to 3PLE, and 743 annotations unique to 3:3:2 (**Figure 2E**). The multiphasic extractions demonstrated high overlap in coverage with three of the next four largest intersections containing a combination of them. Overall, these data demonstrate the efficient phase partitioning of the metabolites with minimal overlap between polar and non-polar phases in all extraction methods. Although the triphasic extraction resulted in more metabolic features, SmartFormula annotations, and identified metabolites, the reproducibility was among the lowest compared to the other extraction methods ranging from 0.9466 to 0.9578 across extraction replicates. Collectively, these data suggest that the Matyash method would be most beneficial in discovery experiments wherein the enrichment of metabolites from the human heart is limited to only one extraction method as it provided a fair number of identified metabolites in addition to extraction reproducibility.

**Figure 2.**
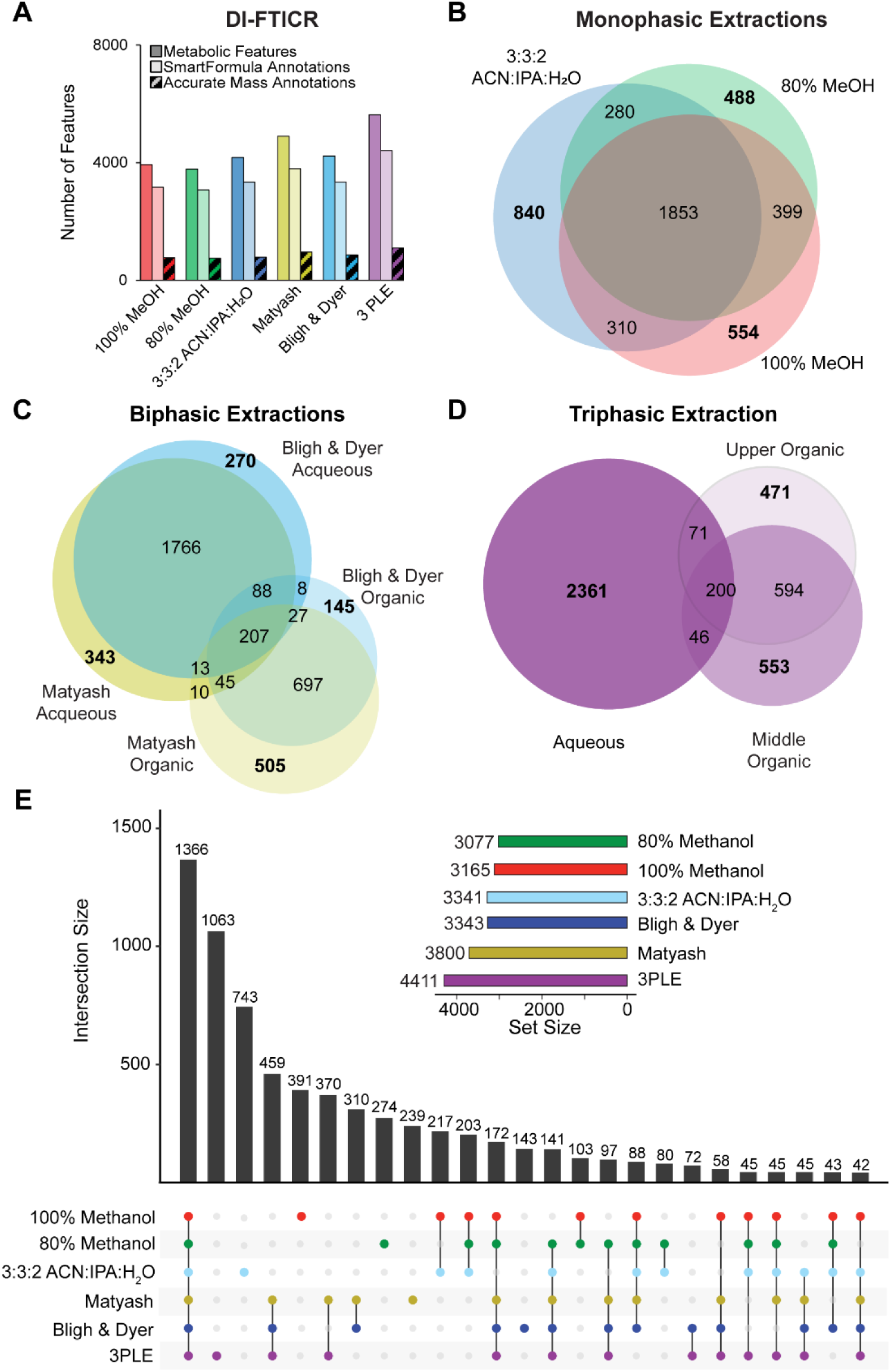
Analysis of human heart metabolites using parallel extractions and ultrahigh-resolution mass spectrometry. Metabolite extracts were analyzed using ultrahigh-resolution DI-FTICR MS. (A) Bar graph comparing the number of metabolic features, SmartFormula annotations (<2.0ppm), and accurate mass annotations (<2.0ppm) across different extraction methods (monophasic (100% MeOH, 80% MeOH and 3:3:2 ACN:IPA:H20), biphasic (Matyash and Bligh and Dyer) and triphasic (3PLE). The number of SmartFormula annotations was compared for the (B) monophasic, (C) biphasic, and (D) triphasic extraction methods. (E) UpSet plot showing the top 25 largest intersections of SmartFormula annotations.

### Complementary MS Platforms for Metabolite Identification

DI-FTICR MS has numerous advantages for metabolite detection and identification. Direct infusion of metabolite extracts bypasses issues of chromatographic compatibility and retention time variation.^33^ Additionally, lower abundant metabolites can be detected due to the spectral summing capability and accurate mass annotations with its high mass accuracy (<2.0 ppm) and isotopic fine structures determination,^34^ which is shown in **Figure 3A-3B**. Additionally, the isotopic distribution was evaluated for selected metabolites (**Figure 3C-3E**, Figures S7-S8). When comparing the mass spectra for fatty acyl esters of hydroxy fatty acids (FAHFA) 40:5, diacylglycerol (DG) 32:0, and N-palmitoyl glutamic acid, we observe impressive alignment between the theoretical and experimental isotopic distributions (**Figure 3C).** Utilizing this platform, we detected 9521 metabolic features, wherein 7,699 and 1,756 could be assigned a chemical formula (SmartFormula annotations) and an accurate mass identification, respectively (**Figure 2A**, Table S1). The drawbacks of the DI-FTICR MS platform are a limited ability for high throughput MS/MS analysis and the separation of isomeric species.

**Figure 3.**
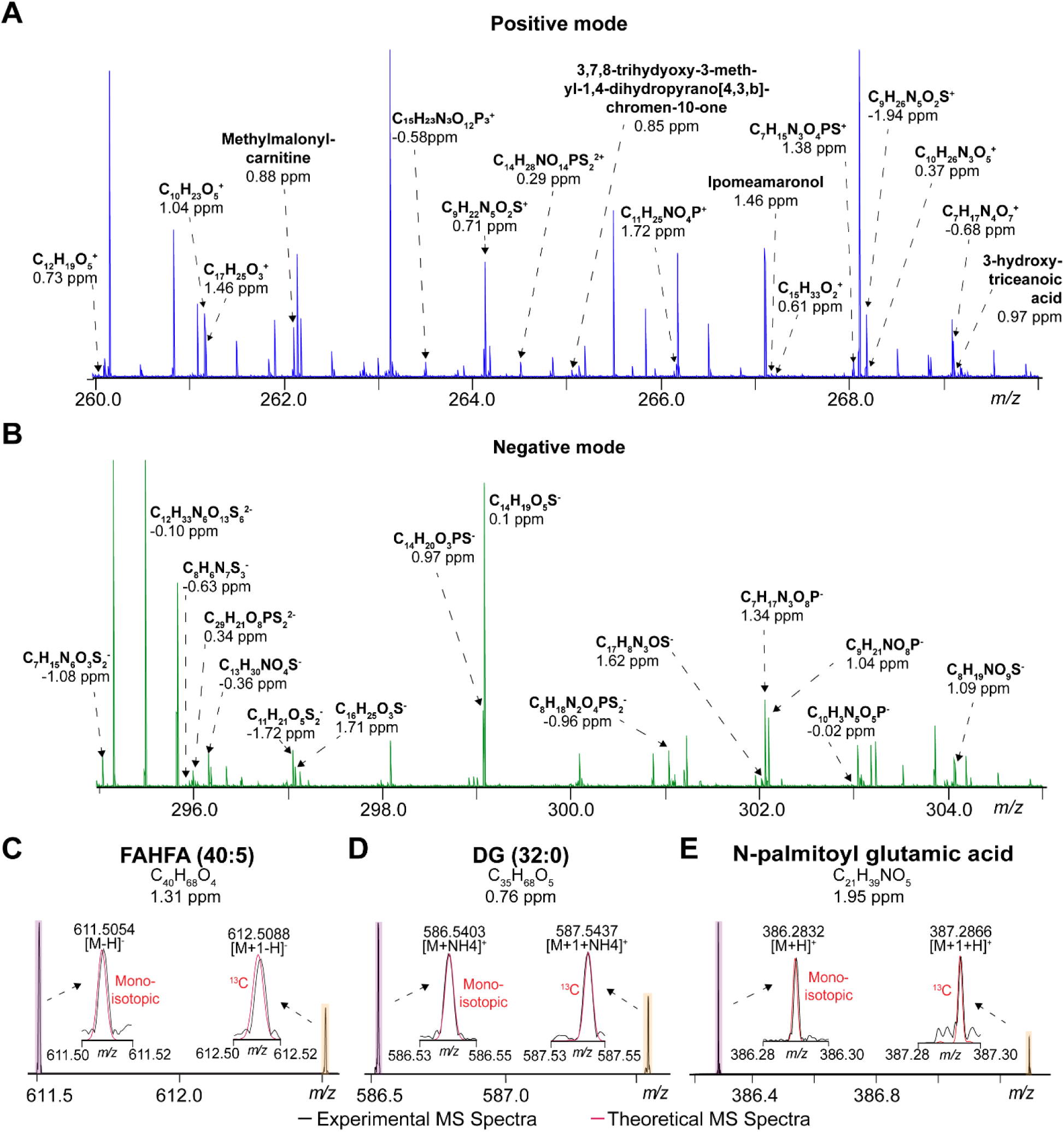
DI-FTICR MS analysis of metabolites in both positive and negative modes. Shown are representative MS spectra (100% MeOH) for (A) positive and (B) negative ion modes for a zoomed *m/z* 10 window where SmartFormula and accurate mass annotations are indicated for each. Experimental mass spectra (black) were overlaid with theoretical mass spectra (red) to compare the isotopic distribution of each. Shown are (C) FAHFA (40:5), (D) DG (32:0), and (E) N-palmitoyl glutamic acid.

High-resolution LC-Q-TOF MS/MS provides complementary means to address these drawbacks.^25^ By searching acquired MS/MS spectra against publicly available libraries, we identified both metabolites (**Figure 4A**) and lipids (**Figure 4B-D**). For lipids, we obtain acyl chain information from the fragmentation patterns by observing the acyl chain directly or by neutral loss of the acyl chain in negative or positive ion modes, respectively (**Figure 4B, 4D**). Of note, two of these lipids, FAHFA 18:1_22:4 and DG 16:0_16:0 match the summed acyl chain composition observed in **Figure 3C-D**, demonstrating the strengths of the LC-Q-TOF-MS/MS platform to gain *sn1* and *sn2* acyl chain information. In the LC-QTOF MS/MS analyses, 21,428 metabolic features were detected, where 626 were identified based on fragmentation matching against publicly available libraries (Table S1). Collectively, our DI-FTICR MS and LC-Q-TOF MS/MS platforms increased the coverage of the human heart metabolome compared to any single MS detection platform alone.

**Figure 4.**
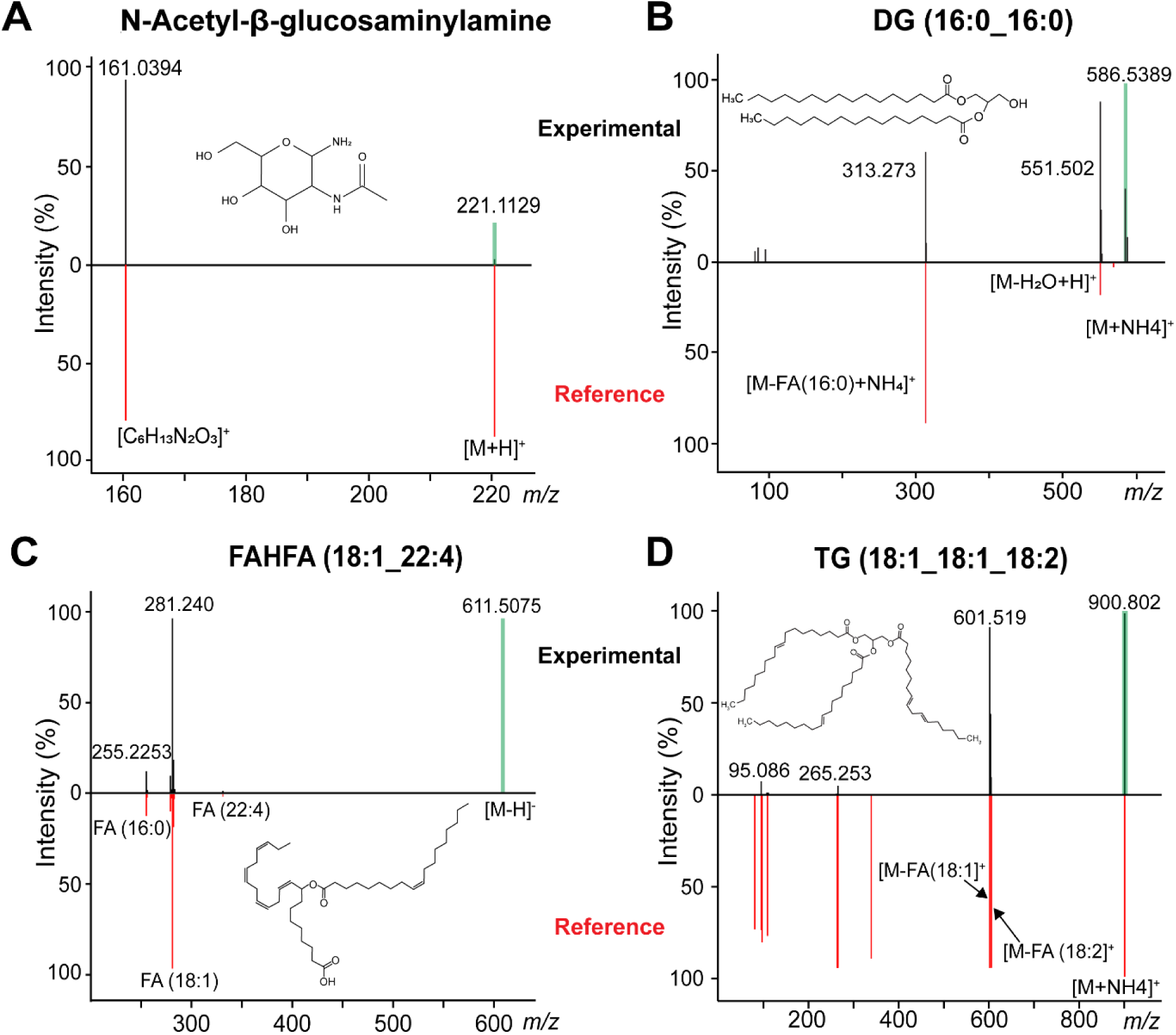
Identification of metabolites using LC-QTOF-MS/MS. Representative LC-QTOF tandem mass spectra were shown. Shown are mirror plots comparing experimental mass spectra (black) with library (reference) mass spectra (red). Peak annotations for (A) N-Acetyl-B-glucosaminylamine, (B) DG (16:0_16:0), (C) FAHFA (18:1_22:4), and (D) TG (18:1_18:1_18:2). FA=fatty acids.

### Identification of Metabolites from Human Heart Tissue

Using multiple extraction and MS detection methods, we identified 2,276 metabolites from human heart tissue (**Figure 5**). More metabolites were identified by accurate mass annotation in the DI-FTICR MS analysis in the multiphasic extractions, however more metabolites were identified with fragmentation matching in the monophasic extractions (**Figure 5A**). Collectively, the triphasic (3PLE) extraction method enriched the most metabolic species when considering chemical class, except for amino acids and glycerophosphates (**Figure 5B**); more amino acids were identified in the Matyash extraction and glycerophosphates in the 100% methanol extraction. Each extraction method contributed uniquely detected metabolites (**Figure 5C**, Table S2); the triphasic (3PLE) extraction provided the greatest at 284, followed 100% MeOH, 3:3:2 ACN:IPA:H2O, and 80% MeOH with 95, 95, and 46 metabolites, respectively. It is important to note that many uniquely detected metabolites in the 100% MeOH extraction were phospholipids, including several phosphatidic acids. In contrast, the 3PLE extracted more lipids across the classes and not one more favorably (Table S2). Markedly, Matyash and B&D contributed the fewest uniquely detected metabolites with 43 and 34, respectively. This observation can likely be explained by the high degree of overlap between the triphasic and biphasic extractions (**Figure 3E**). By utilizing parallel extractions, we detected an additional 800 metabolites compared to the best performing single extraction of the human heart alone.

**Figure 5.**
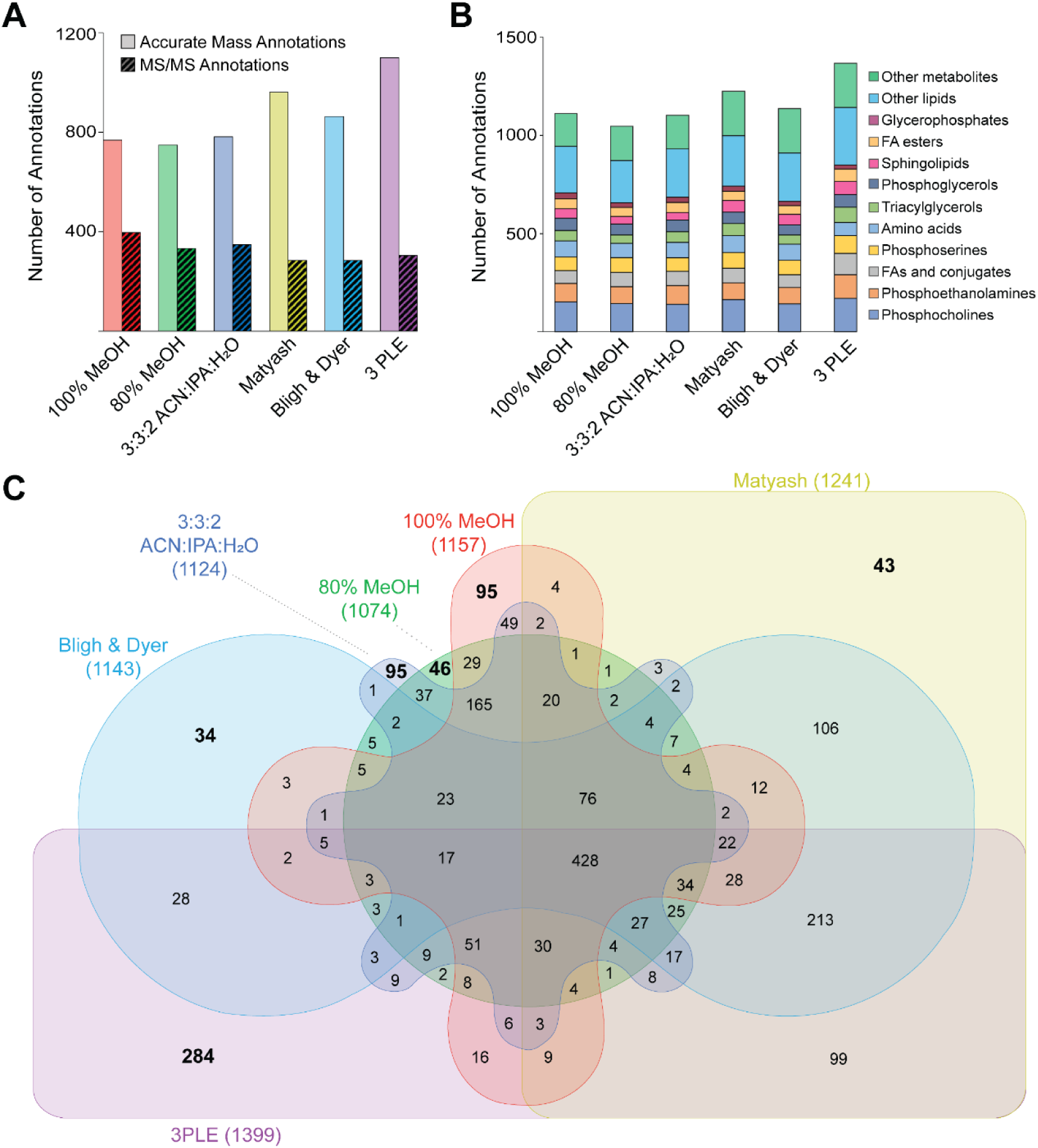
Summary of identified human heart metabolites. Metabolite extracts were analyzed by DI-FTICR and LC-Q-TOF-MS/MS. For LC-Q-TOF MS/MS analyses, monophasic extracts were analyzed using both HILIC and RP chromatography and aqueous and organic biphasic and triphasic extracts were analyzed using HILIC and RP chromatography, respectively. (A) Bar graph showing accurate mass annotations (DI-FTICR accurate mass annotations (≤2.0ppm) and LC-Q-TOF MS/MS spectral library matches (≤20.0ppm, MetaboScape MS/MS score ≥400)) across different extraction methods (monophasic (100% MeOH, 80% MeOH and 3:3:2 ACN:IPA:H2O), biphasic (Matyash and Bligh and Dyer), and triphasic (3 phase liquid extraction, 3PLE). (B) Bar graph showing the top 10 molecular classes of combined annotations by extraction. (C) Venn diagram showing the number of non-overlapping (bolded) combined annotations detected in each extraction method.

Further characterization including compound classification and pathway analysis was performed on all metabolites (**Figure 6**, Table S1). In the compound classification analysis, many compounds were observed to be *Lipids and lipid-like molecules*, the majority of which were *glycerophosphocholines (PCs)*, *glycerophosphoethanolamines (PEs)*, and *fatty acids and conjugates* with 152, 94, and 65 metabolites, respectively (**Figure 6A**). The non-lipid compounds were largely *Organic acids and derivates* containing 209 metabolites, the majority of which 146 were *Amino acids, peptides, and analogues.* To gain more insight regarding the polarity of metabolites that were enriched by our approach, we performed a ChemRich analysis (Figure S6). This analysis revealed a broad range of metabolites, spanning the entire polarity spectrum. Similar to the compound classification analysis (**Figure 6A**), we observed a larger mapping of *Lipid and lipid-like molecules*. Additionally, pathway analysis was performed to elucidate biological relevance of all metabolites (**Figure 6B**). Among the enriched pathways was Purine Metabolism, Glycerophospholipid Metabolism, and Linoleic acid Metabolism, all key pathways involved in metabolic processes of the human heart.^35–38^ To illustrate the enhanced coverage of the metabolome by using our approach, we selected a pathway known to be altered in many cardiac diseases, purine metabolism,^8, 12–14^ and highlighted the extraction method(s) in which each metabolite was found (**Figure 6C**). Remarkably, four of the metabolites were observed in only a single extraction - ribose-5-phosphate (R5P), guanine diphosphate (GDP), inosine diphosphate (IDP), and cyclic adenosine monophosphate (cAMP). Many metabolites including glutamine, adenosine, adenosine diphosphate (ADP), and hypoxanthine were observed in all six extraction methods. Moreover, B&D was observed to enrich the most metabolites involved in this pathway; 3PLE was observed to enrich the least, despite its ability to enrich the largest number of metabolites. Further, 5-Aminoimidazole-4-carboxamide ribonucleotide (AICAR) and guanidine triphosphate (GTP) were observed only in the monophasic extractions. Together, these data provide critical insight regarding the selection of an appropriate extraction method(s) for the analysis of human heart metabolites.

**Figure 6.**
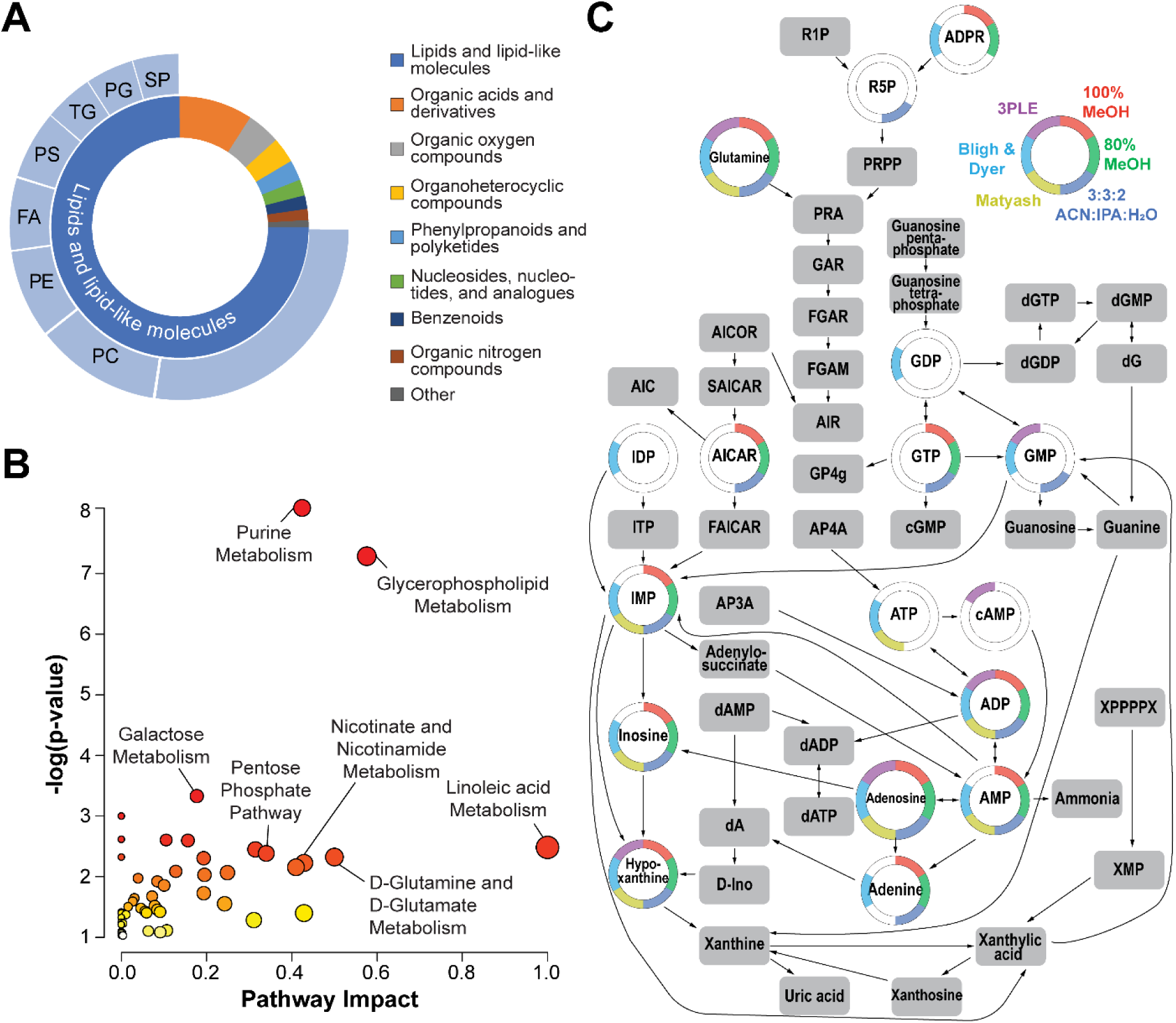
Molecular classification and pathway analysis of identified human heart metabolites. (A) Double-donut plot showing the most enriched metabolite classes. (B) Summary of pathway analysis for all identified metabolites where the colors and size of each circle is based on *p-*value and pathway impact value (high (red) to low (yellow)), respectively. (C) Shown is the pathway for purine metabolism. Increased pathway coverage is achieved by using parallel extraction methods compared to any single extraction method alone. Color indicates the extraction method from which the metabolite was identified.

### Characterization of Human Heart Metabolome

In this study, we combined parallel extractions and complementary MS detection platforms to maximize the human heart metabolome coverage. **Table 1** shows a summary of metabolites that are known to be implicated in heart diseases as previously reported.^7–14^ Carnitine, choline, and ACars (various chain lengths) were broadly observed across all extraction methods and MS-based detection platforms. Amino acids were observed across most extractions; however, detection was observed to be better suited on the LC-QTOF MS/MS platform. Of note, amino acids and ACars have been reported to return to comparable levels of healthy controls after left ventricle assist device (LVAD) surgery.^9^ ATP was not observed in any of the monophasic extraction methods and detection was observed to be better suited on the DI-FTICR MS platform. Previously, Previs and co-authors noted ATP had an inverse correlation with the size of the left atria in hypertrophic cardiomyopathy (HCM).^13^ Both NAD and S-Adenosylmethionine were observed in nearly half of the extractions and both MS-based platforms. LPC 18:2 was only enriched in the 100% MeOH and 3PLE extraction methods and only identified by LC-QTOF MS/MS. CE 18:1, which is associated with cardiovascular disease risk,^39^ was not observed in the Matyash or B&D extractions and only identified by LC-QTOF MS/MS. Detecting this broad range of metabolites is essential as optimizing energy substrate usage is a main function of the heart.^40–43^

**Table 1.**
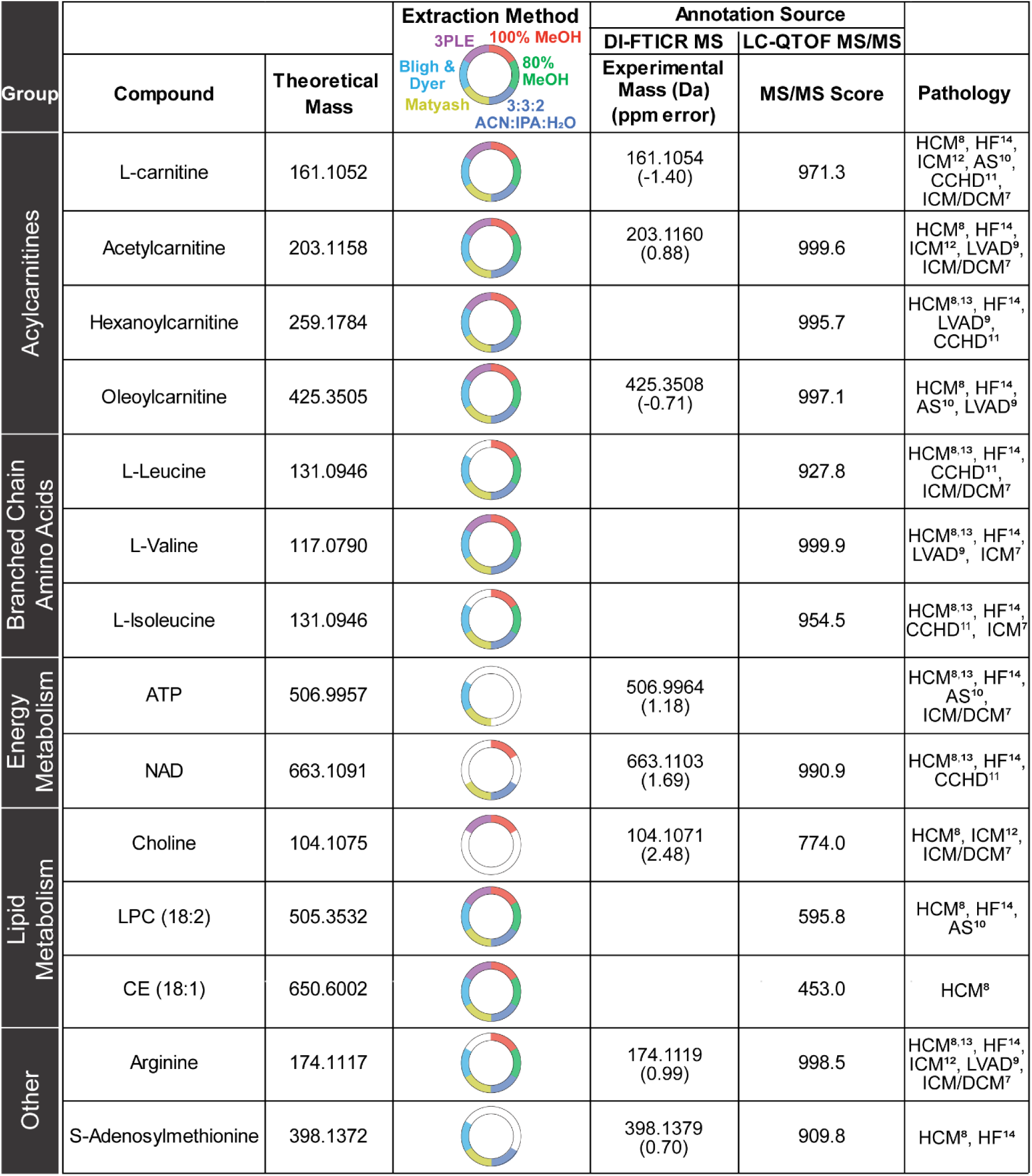
Summary of identified critical cardiac metabolites which are implicated in heart diseases. Summary of identified metabolites which overlap with previous human heart analyses. Representative metabolites shown for each group where extraction and MS detection platform are indicated. Metabolites detected by FT-ICR were ≤2.0 ppm mass error. Metabolites detected by LC-QTOF-MS/MS have a MS/MS score ≥ 400, a threshold which was implemented in this study based on manual inspection of all acquired MS/MS spectra. HCM=Hypertrophic cardiomyopathy, HF=heart failure, AS=aortic stenosis, CCHD=cyanotic congenital heart disease, ICM=ischemic cardiomyopathy, DCM=dilated cardiomyopathy, and LVAD=left ventricular assist device.

| Group | Compound | Theoretical Mass | Extraction Method<br>3PLE 100% MeOH<br>Bligh & Dyer<br>Matyash<br>80% MeOH<br>3:3:2<br>ACN:IPA:H<sub>2</sub>O | Annotation Source |  | Pathology |
| --- | --- | --- | --- | --- | --- | --- |
|  |  |  |  | DI-FTICR MS<br>Experimental Mass (Da)<br>(ppm error) | LC-QTOF MS/MS<br>MS/MS Score |  |
| Acylcarnitines | L-carnitine | 161.1052 |  | 161.1054<br>(-1.40) | 971.3 | HCM<sup>8</sup>, HF<sup>14</sup>,<br>ICM<sup>12</sup>, AS<sup>10</sup>,<br>CCHD<sup>11</sup>,<br>ICM/DCM<sup>7</sup> |
|  | Acetylcarnitine | 203.1158 |  | 203.1160<br>(0.88) | 999.6 | HCM<sup>8</sup>, HF<sup>14</sup>,<br>ICM<sup>12</sup>, LVAD<sup>9</sup>,<br>ICM/DCM<sup>7</sup> |
|  | Hexanoylcarnitine | 259.1784 |  |  | 995.7 | HCM<sup>8,13</sup>, HF<sup>14</sup>,<br>LVAD<sup>9</sup>,<br>CCHD<sup>11</sup> |
|  | Oleoylecarnitine | 425.3505 |  | 425.3508<br>(-0.71) | 997.1 | HCM<sup>8</sup>, HF<sup>14</sup>,<br>AS<sup>10</sup>, LVAD<sup>9</sup> |
| Branched Chain Amino Acids | L-Leucine | 131.0946 |  |  | 927.8 | HCM<sup>8,13</sup>, HF<sup>14</sup>,<br>CCHD<sup>11</sup>,<br>ICM/DCM<sup>7</sup> |
|  | L-Valine | 117.0790 |  |  | 999.9 | HCM<sup>8,13</sup>, HF<sup>14</sup>,<br>LVAD<sup>9</sup>, ICM<sup>7</sup> |
|  | L-Isoleucine | 131.0946 |  |  | 954.5 | HCM<sup>8,13</sup>, HF<sup>14</sup>,<br>CCHD<sup>11</sup>, ICM<sup>7</sup> |
| Energy Metabolism | ATP | 506.9957 |  | 506.9964<br>(1.18) |  | HCM<sup>8,13</sup>, HF<sup>14</sup>,<br>AS<sup>10</sup>,<br>ICM/DCM<sup>7</sup> |
|  | NAD | 663.1091 |  | 663.1103<br>(1.69) | 990.9 | HCM<sup>8,13</sup>, HF<sup>14</sup>,<br>CCHD<sup>11</sup> |
| Lipid Metabolism | Choline | 104.1075 |  | 104.1071<br>(2.48) | 774.0 | HCM<sup>8</sup>, ICM<sup>12</sup>,<br>ICM/DCM<sup>7</sup> |
|  | LPC (18:2) | 505.3532 |  |  | 595.8 | HCM<sup>8</sup>, HF<sup>14</sup>,<br>AS<sup>10</sup> |
|  | CE (18:1) | 650.6002 |  |  | 453.0 | HCM<sup>8</sup> |
| Other | Arginine | 174.1117 |  | 174.1119<br>(0.99) | 998.5 | HCM<sup>8,13</sup>, HF<sup>14</sup>,<br>ICM<sup>12</sup>, LVAD<sup>9</sup>,<br>ICM/DCM<sup>7</sup> |
|  | S-Adenosylmethionine | 398.1372 |  | 398.1379<br>(0.70) | 909.8 | HCM<sup>8</sup>, HF<sup>14</sup> |

New metabolites identified using our combined parallel extraction and complementary MS-detection approach are summarized in **Table 2**. The isotopic distribution (DI-FTICR analysis) and fragmentation matching (LC-QTOF-MS/MS analysis) are shown for a subset of these (Figures S7-S10). Overall, a growing amount of evidence suggests that many of these metabolites may be of interest in understanding cardiac metabolism.^44–49^ Previous reports indicate that FAHFAs are involved in the regulation of glucose metabolism and inflammation;^44^ in this study, we show detection of FAHFA 40:5. Other reports implicate DGs in the activation of protein kinase C and D isoforms and subsequent regulation of glucose and lipid metabolism; ^44,45, 46^ we observed DG 16:0/16:0 in our comprehensive metabolomic analysis. Markedly, FAHFA (40:5) was observed in most extractions except B&D and DG 16:0/16:0 was only observed in the 3PLE extraction. Ceramides and their precursors, sphingomyelins, are involved in lipotoxicity.^47, 48^ Moreover, ceramides and sphingomyelins with longer acyl chains, such as N-lignoceroyslsphingosine, have been associated with better cardiovascular disease outcomes.^47^ On the other hand, ceramides and sphingomyelins with shorter chains, such as N-stearoyl-erythro-sphingosine, have been associated with poorer outcomes.^47^ Further, reducing total ceramide levels in rodent models revealed decreased onset of cardiovascular complications.^48^ Both ceramides and sphingomyelins were detected by LC-QTOF MS/MS with N-stearoyl-erythro-sphingosine observed in four liquid-liquid extractions and N-lignoceroyslsphingosine observed only in the 100% MeOH extraction. The carbohydrate conjugates, β-L-fucose-1-phosphate and N-acetyl-β-glucosaminylamine, were observed by LC-Q-TOF MS/MS in most metabolite extractions. Interestingly, these are protein glycosylation pathway products which have been observed to be dysregulated in streptozotocin-induced mouse models of diabetic cardiomyopathy,^50,49^ however, they have not been reported in the human heart studies. The identification of these new metabolites may enhance our understanding of cardiac metabolism.

**Table 2.**
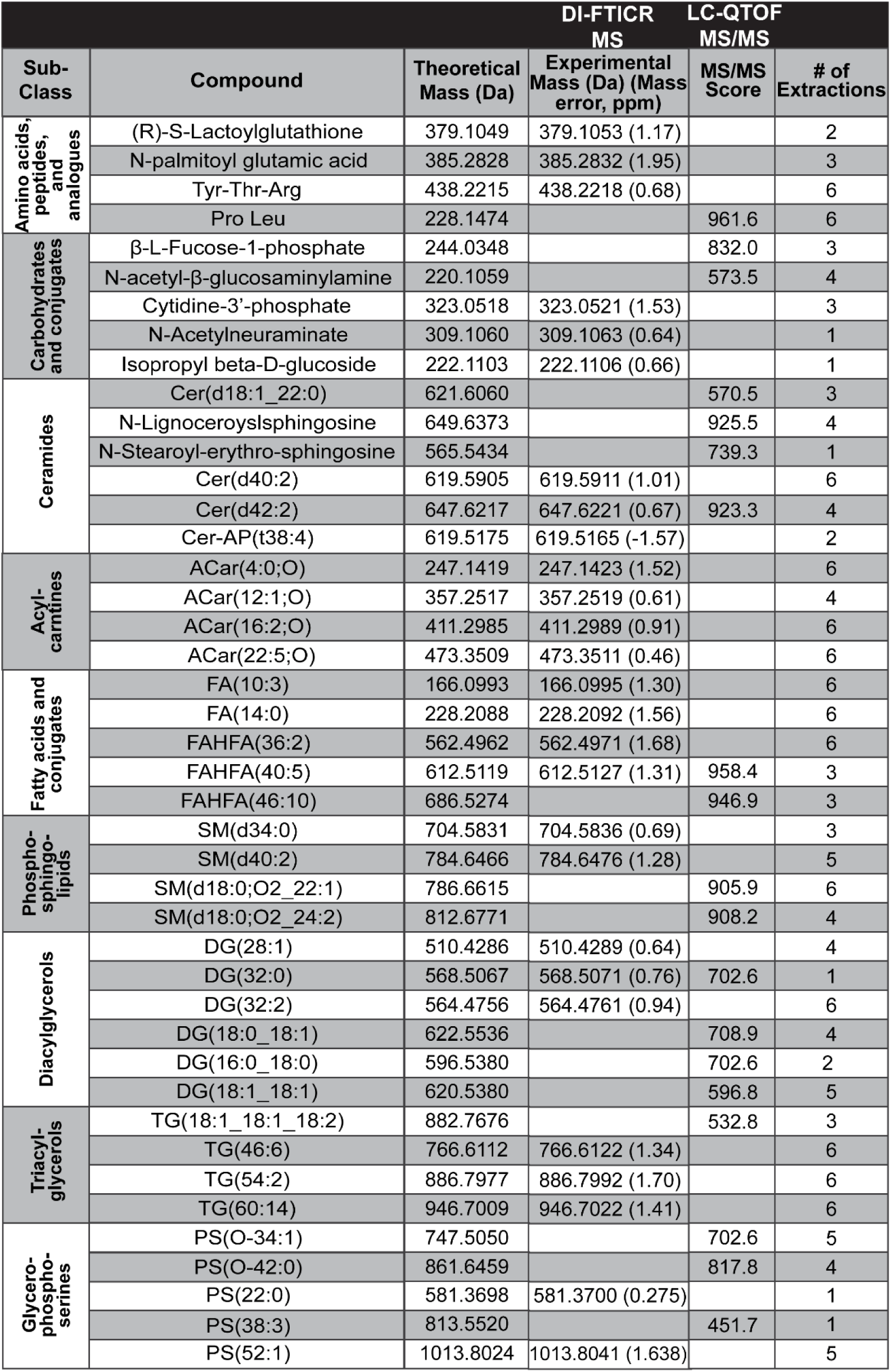
Summary of newly identified metabolites in the cardiac metabolome. Table summarizing metabolites grouped by subclass which are uniquely identified by using parallel extractions and complementary MS detection methods. Metabolites detected by FT-ICR were ≤2.0 ppm mass error. Metabolites detected by LC-QTOF-MS/MS have a MS/MS score ≥ 400.

## Conclusions

Herein, we describe a comprehensive analysis of the human heart metabolome where six parallel liquid-liquid extraction methods (three monophasic, two biphasic, and one triphasic) and two complementary MS-based detection platforms (DI-FTICR and LC-QTOF MS/MS) were utilized for the comprehensive metabolomic analysis. Using this approach, 2276 metabolites were identified from healthy, human donor heart tissue, and represent the deepest coverage of the human heart metabolome to date. Overall, the triphasic extraction was observed to yield nearly 1400 metabolites and best suited for enriching lipids; however, the number of polar metabolites in this extraction was much less compared to the biphasic, Matyash extraction. The triphasic extraction method was observed to be the least reproducible of the six considered. All of the monophasic extractions were observed to perform similarly with 1074-1157 metabolites identified where the 3:3:2 ACN:IPA:H_2_0 extraction was observed to be the most reproducible. With this approach, we identified new metabolites in the human heart. We envision that the information in this study may be useful in selecting an appropriate extraction and MS detection platform to assist in metabolomic analyses of the human heart.

## Data Availability

All data contained within this manuscript are available upon reasonable request of the corresponding author. All mass spectrometry data have been deposited to MASSive with the dataset identifier MSV000091539.

## Acknowledgements

This research is supported by NIH R01 HL109810-06 (Y.G.). Y.G. acknowledges NIH R01 GM125085, R01 HL096971, and S10 OD018475. B.W. acknowledges the University of Wisconsin Department of Cell and Regenerative Biology, Department of Pathology and Laboratory Medicine, NHLBI T32 HL007936, and NIH F31 HL152647 for funding and use of its facilities and services.

