## Supporting Information for "Comprehensive Metabolomic Analysis of Human Heart Tissue Enabled by Parallel Metabolite Extraction and High-Resolution Mass Spectrometry"

Ying Ge^1,2,^*

**^1^** Department of Cell and Regenerative Biology, University of Wisconsin-Madison, Madison, Wisconsin, 53705, USA

^2^ Human Proteomics Program, School of Medicine and Public Health, University of Wisconsin-Madison, Madison, Wisconsin, 53705, USA

^†^ Denotes equal contribution

*To whom correspondence should be addressed:

Professor Ying Ge Ph.D.

1111 Highland Ave, WIMR II 8551

Madison, Wisconsin 53705, USA

Or

Melissa R. Pergande Ph.D.

1111 Highland Ave, WIMR II 8546

Madison, Wisconsin 53705, USA

**Experimental Section**

*DI-FTICR MS analysis of metabolite extracts*

Fourier transform ion cyclotron resonance mass spectrometer (FTICR MS) detection of metabolites was performed using direct infusion (DI) by syringe pump into a Bruker solariX 12T FTICR mass spectrometer (Bruker Daltonics, Bremen, Germany). For metabolite resuspension, monophasic and polar phase extracts (upper phase from B&D, lower phase from Matyash, and lower phase from 3PLE) were resuspended in 300 µL of 50:50 methanol: water, while nonpolar phases (lower phase from B&D, upper phase from Matyash, and middle and upper phase from 3PLE) were resuspended in 300 µL of 90:10 isopropanol: acetonitrile. Resuspension modifiers were 10 mM ammonium formate and 0.1 % formic acid for positive mode and 10 mM ammonium acetate for negative mode. Samples were drawn into a syringe and directly infused at 3 µL/min via a 100 µm x 40 cm PEEK tubing into the FTICR MS ion source. Ions were accumulated for 0.1 s, and an 8M transient size applied with 300 scans collected. The *m/z* range was set to 40-1500, with *m/z* 50 Q1 mass. Dry gas flow was set to 4L/min at 150 ºC. To improve ion transition, the largest frequency values for octupole (5 MHz), quadrupole (2 MHz), and transfer hexapole (6 MHz) were used. Time of flight was set to 0.8 ms. Sweep excitation power was set to 27%. The estimated resolving power at *m/z* 400 was 190,000. The FTICR MS was calibrated with Agilent ESI-L tuning mix in both positive and negative modes before experiments. The syringe was cleaned with 100% MeOH between samples.

FTICR mass spectra were processed using ftmsProcessing software V2.3.0 (Bruker Daltonics, Bremen, Germany) to remove Gibbs and harmonic peaks. The resulting data files were analyzed using MetaboScape v2022b (Bruker Daltonics, Bremen, Germany). Bucket (mass) lists in positive and negative ion modes were generated using the T-ReX 2D workflow. The maximum mzDelta was set to 2 mDa, maximum charge state was set to 2, and the intensity threshold was set to 5E7. The minimum number of features for the result was set to 2 of 3 replicates per phase. Bucket lists in positive and negative ion modes were merged into one with 1.0 ppm *m/z* tolerance. The merged bucket list was annotated with the SmartFormula function with 1.0 ppm as the narrow Δ*m/z* cutoff, 2.0 ppm as the wide Δ*m/z* cutoff, 20 as the narrow mSigma cutoff, and 50 as the wide mSigma cutoff. Putative metabolites were annotated by target lists generated from MassBank of North America database with a 2.0 ppm mass error cutoff. The annotation search used “hierarchical search”, which selects annotation from first library hit, and “attempt primary ion reassignment” was performed. Preference for target list containing previously detected metabolites with our platform, then outside libraries were considered-LipidBlast, internal MS/MS libraries, and Massbank of North America (MoNA) export for accurate mass matching.

*LC-QTOF-MS/MS analysis of metabolite extracts*

Metabolite extracts from the monophasic, biphasic polar, and triphasic polar lipid extractions were resolved using reversed phase chromatography on a Waters HSS C18 T3 (300 µm x 100 mm, 100 Å, 1.8 µm) column. For positive ion mode, mobile phase A was 0.1% formic acid and mobile phase B (MPB) was MeOH containing 0.1% formic acid. For negative ion mode, mobile phase A was 10 mM ammonium acetate and MPB was MeOH containing 10 mM ammonium acetate. The LC gradient started at 5% MPB, held for 3 min, went to 99% MPB by 8min and held for 9 min, and finally a 3 min re-equilibration. The resuspension solvent and flow rate were 50 µL 95:5 H_2_O: MeOH and 8 µL/min, respectively.

Metabolite extracts from the monophasic, biphasic non-polar, and triphasic polar and non-polar lipid extractions were resolved using reversed phase chromatography on a Waters HSS C18 T3 (300 µm x 100 mm, 100 Å, 1.8 µm) column. Mobile phase A was 3:2 ACN:H2O and mobile phase B was 9:1 IPA:ACN with both having either 10 mM ammonium formate or 10 mM ammonium acetate for positive or negative ion mode, respectively. The LC gradient started at 70% B for 1 min, went to 82% B by 3 min, 99% B by 13 min and held for 4 min, and followed by a 3 min re-equilibration. The resuspension solvent and flow rate were 50 µL 9:1 MeOH: CHCl_3_ and 8µL/min, respectively.

Metabolite extracts from the biphasic polar and triphasic polar phases were resolved using hydrophilic interaction liquid chromatography (HILIC) on a Waters ACQUITY UPLC BEH Amide (1.0 mm x 150 mm, 1.8 µm) column. Mobile phase A was H_2_O and mobile phase B was 95:5 ACN: H2O, both containing 10 mM ammonium formate and 0.125% formic acid in positive ion mode and 10 mM ammonium acetate in negative ion mode. The LC gradient started at 99% B for 2 min, 70% B at 8 min, 40% B at 11.5 min, and 30% B at 14 min, followed by a 6 min re-equilibration. The resuspension solvent and flow rate were 50 µL 80:20 ACN: H_2_O and 12 µL/min, respectively.

**Supplementary Figures**
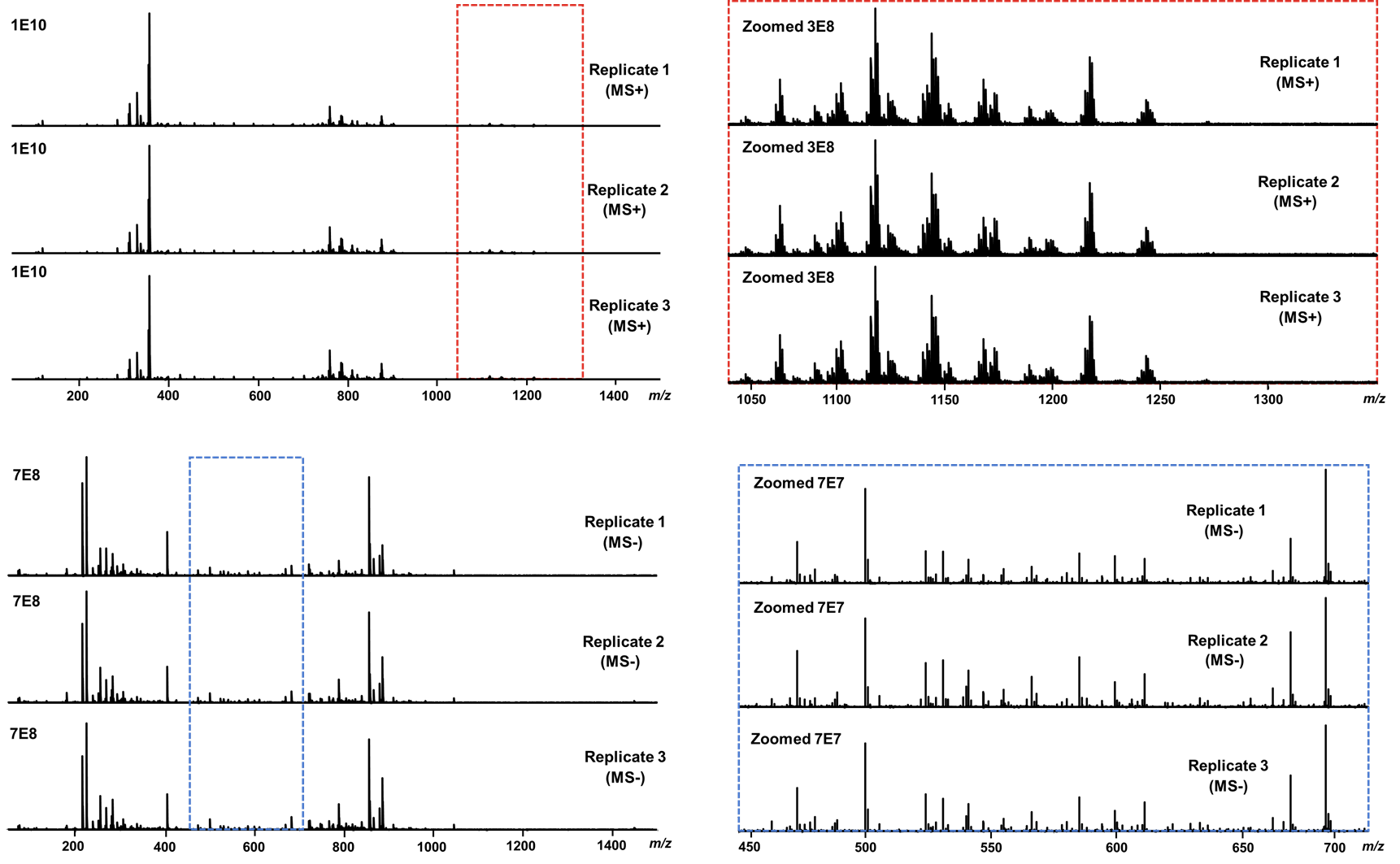

**Figure S1. Representative ultrahigh-resolution mass spectra.** Representative DI-FTICR (Bruker SolariX 12T) mass spectra of triplicate 100% methanol extractions. Shown is the mass spectra for positive (MS+, top) and negative (MS-, bottom) ion modes. Corresponding zoomed regions of low abundance ions are shown to the right.

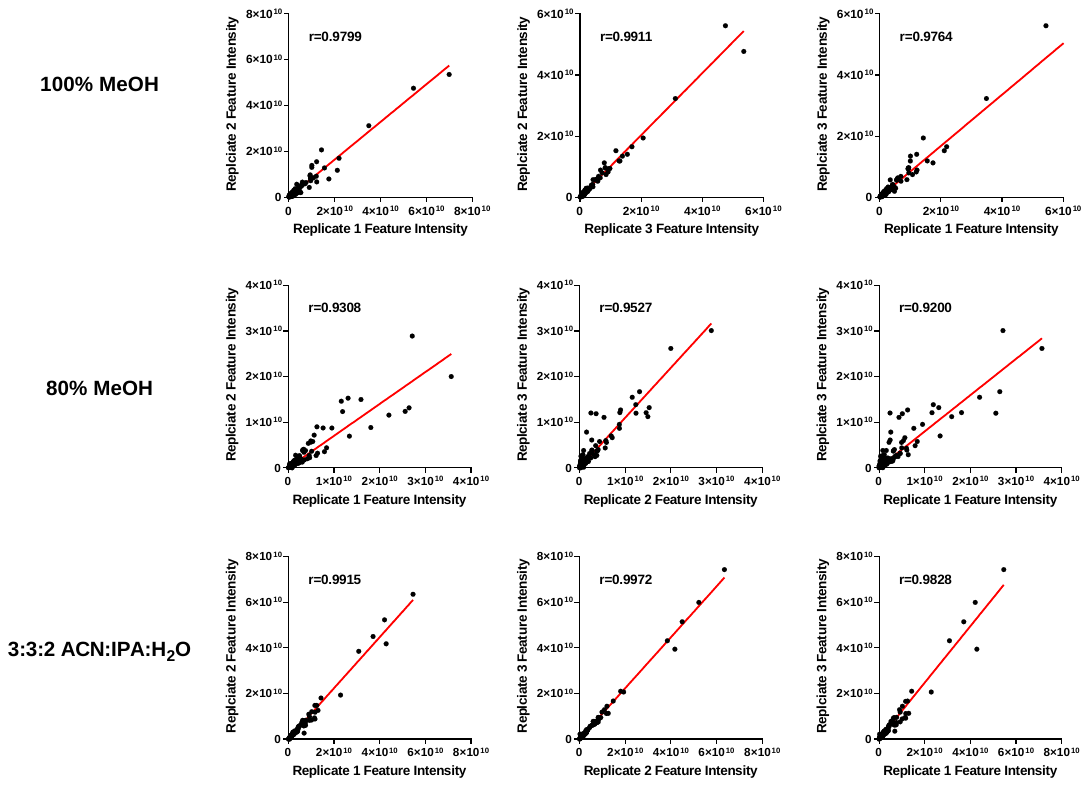

**Figure S2. Evaluation of monophasic extraction reproducibility.** Extraction reproducibility was evaluated across technical replicates for all monophasic (100% MeOH, 80% MeOH and 3:3:2 ACN: IPA:H_2_O) extractions. A Pearson’s correlation coefficient was used to measure the degree of variance between replicates. MeOH = methanol, ACN= acetonitrile, IPA= 2-propanol, and H_2_O= water. Data was collected using ultrahigh-resolution mass spectrometry (DI-FTICR, Bruker SolariX 12T).

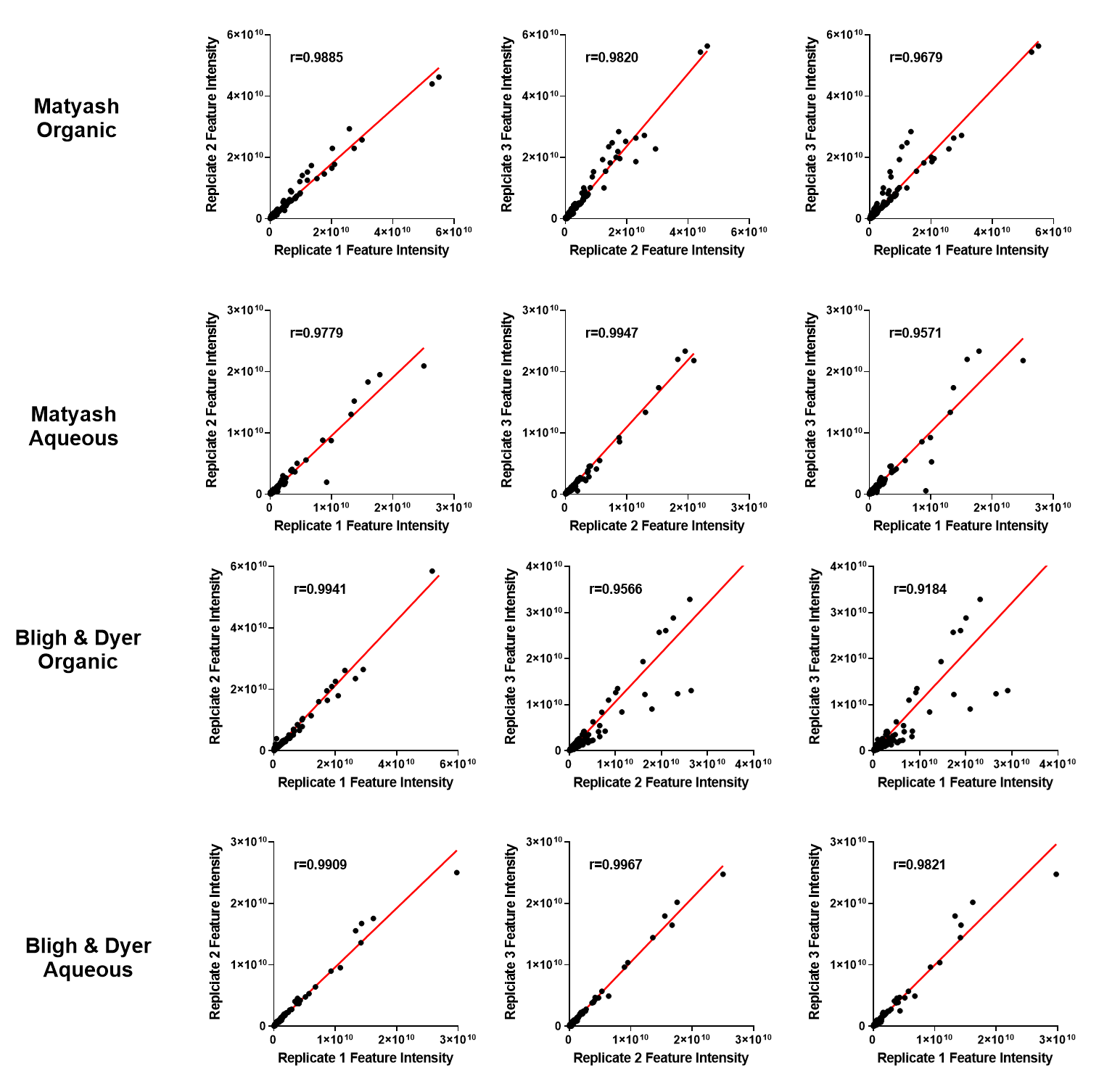

**Figure S3. Evaluation of biphasic extraction reproducibility.** Extraction reproducibility was evaluated across technical replicates for both biphasic extractions (Matyash and Bligh & Dyer). A Pearson’s correlation coefficient was used to measure the degree of variance between replicates. Data was collected using ultrahigh-resolution mass spectrometry (DI-FTICR, Bruker SolariX 12T).

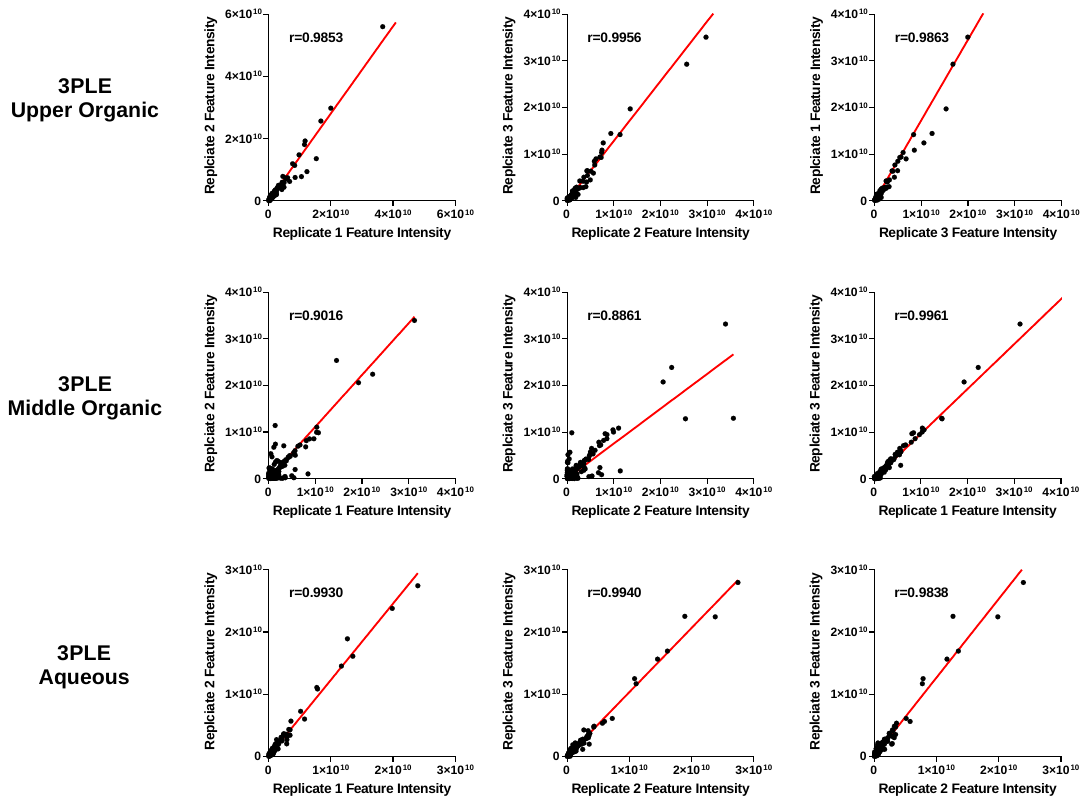

**Figure S4. Evaluation of triphasic extraction reproducibility.** Extraction reproducibility was evaluated across technical replicates for the triphasic extraction (three-phase liquid extraction, 3-PLE) comparing the non-polar (upper organic), polar lipids (middle organic) and non-polar (aqueous) phases. A Pearson’s correlation coefficient was used to measure the degree of variance between replicates. Data was collected using ultrahigh-resolution mass spectrometry (DI-FTICR, Bruker SolariX 12T).

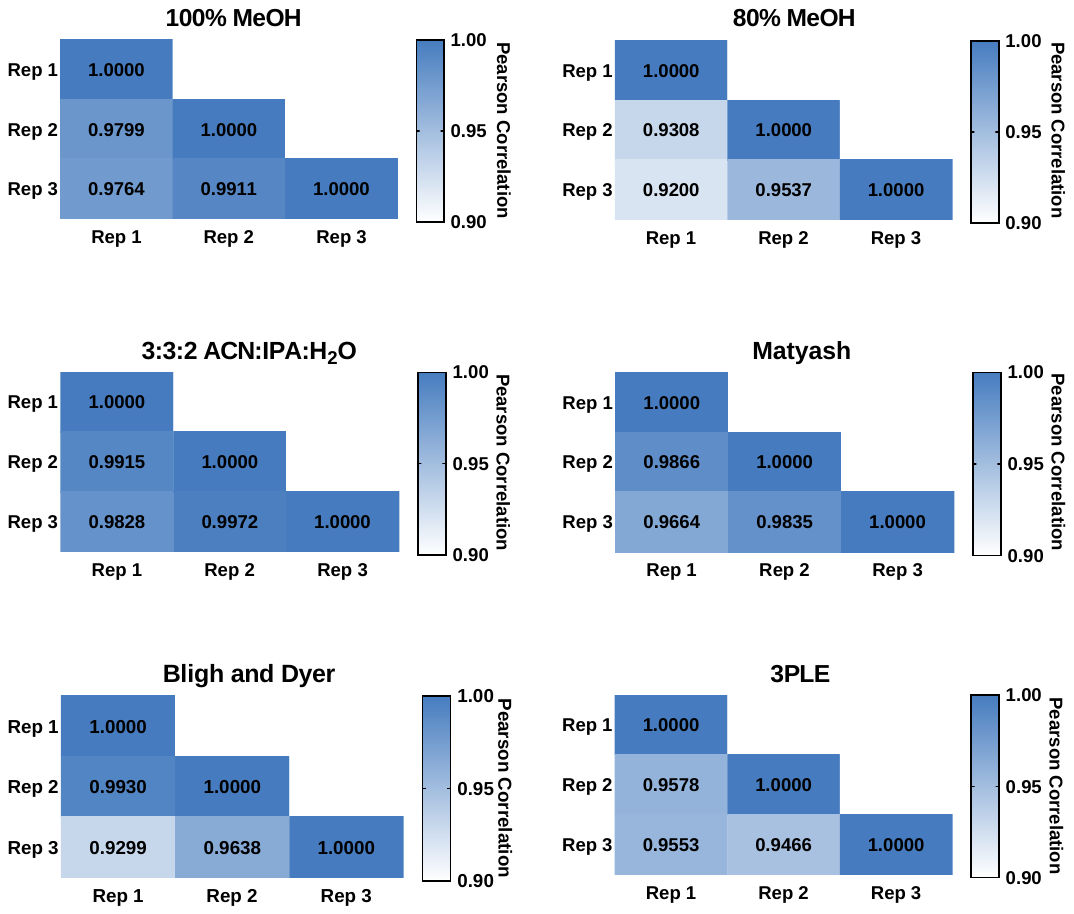

**Figure S5. Comparison of extraction reproducibility across all extraction.** The reproducibility of extractions was evaluated across technical replicates for the monophasic extractions (100% MeOH, 80% MeOH and 3:3:2 ACN:IPA:H_2_O), biphasic extractions (Matyash and Bligh & Dyer), and triphasic extraction (three-phase liquid extraction, 3-PLE) was compared. A Pearson’s correlation coefficient was used to measure the degree of variance between replicates. MeOH = methanol, ACN= acetonitrile, IPA= 2-propanol, and H_2_O= water. Data was collected using ultrahigh-resolution mass spectrometry (DI-FTICR, Bruker SolariX 12T).

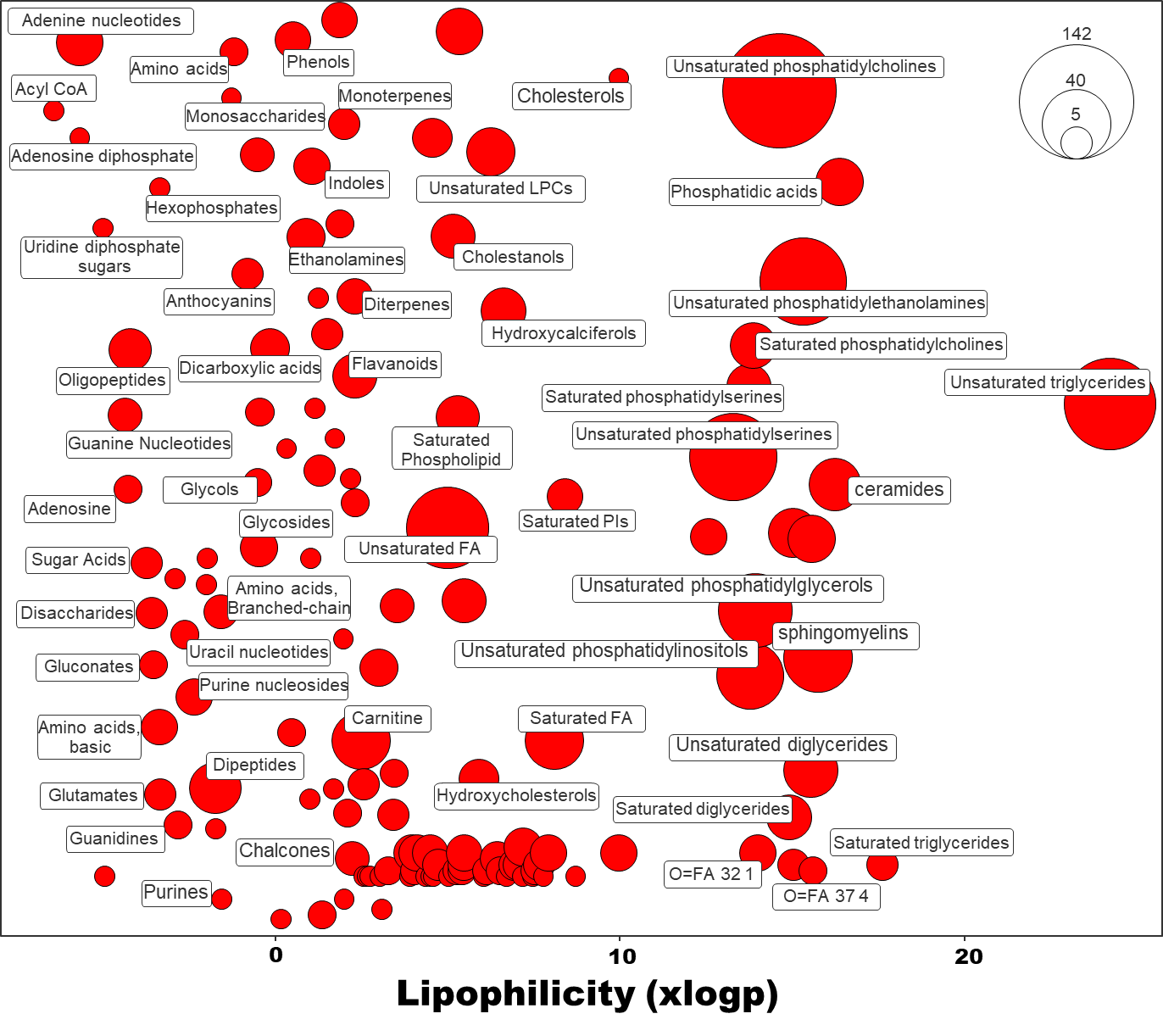

**Figure S6. Chemical similarity enrichment analysis of human heart metabolites.** Combined annotations (Figure 6A) were subject to lipophilicity analysis to visualize the broad range of metabolites detected using multiple extraction methods. ChemRich plot showing the lipophicility (octanol-water partition coefficient, logp) of extracted metabolites. Clusters were based on MeSH (medical subject heading) annotation. The size of each circle indicates the number of metabolites per group.

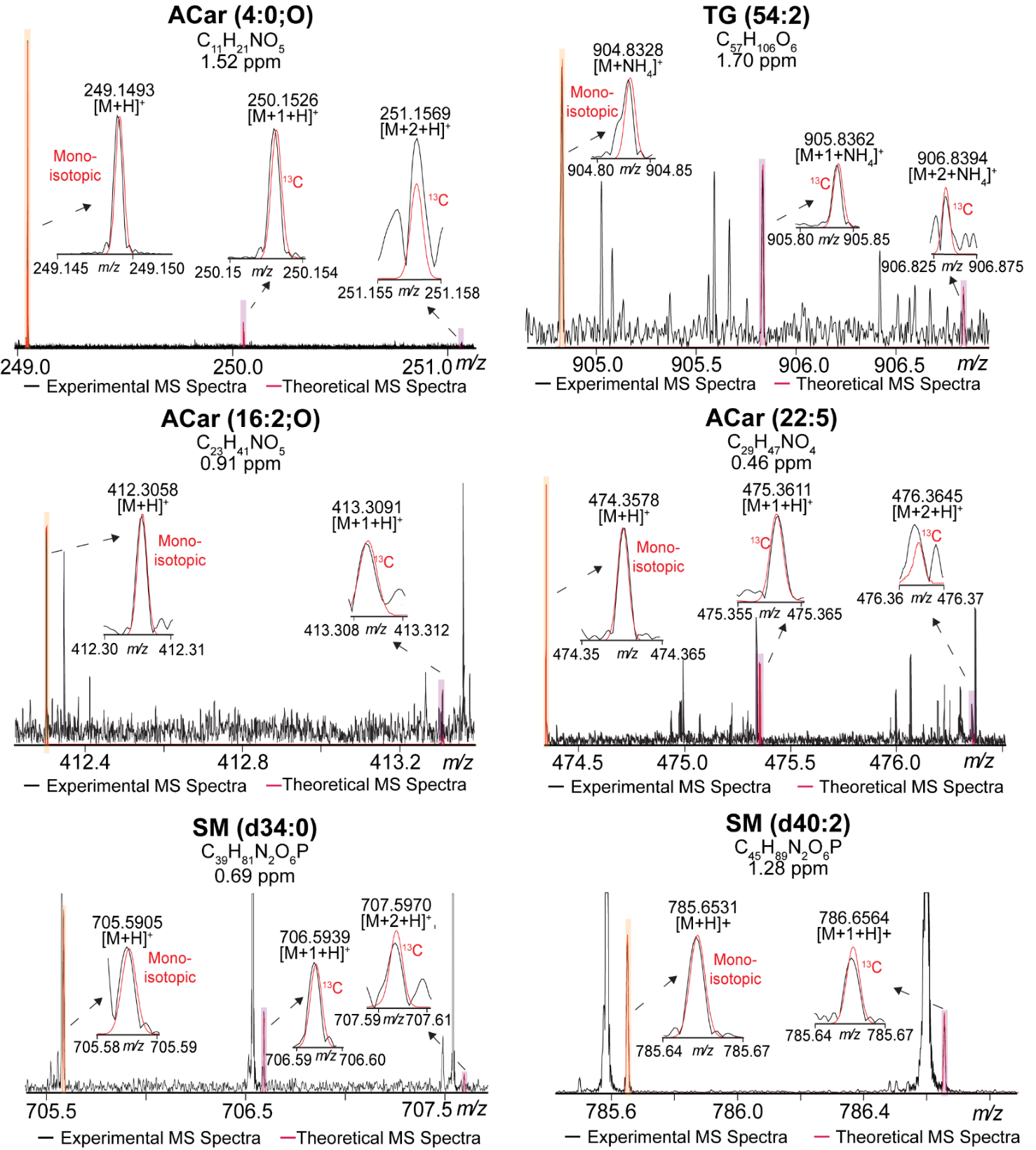

**Figure S7. Representative isotopic fine structure of newly reported metabolites in positive ion mode.** Shown are representative ultrahigh resolution DI-FTICR MS spectra in positive ion mode. Experimental mass spectra (black) were overlaid with theoretical mass spectra (red) to compare the isotopic distribution of newly reported metabolites. ACar=acylcarnitine, TG=triacylglyceride, and SM =sphingomyelin.

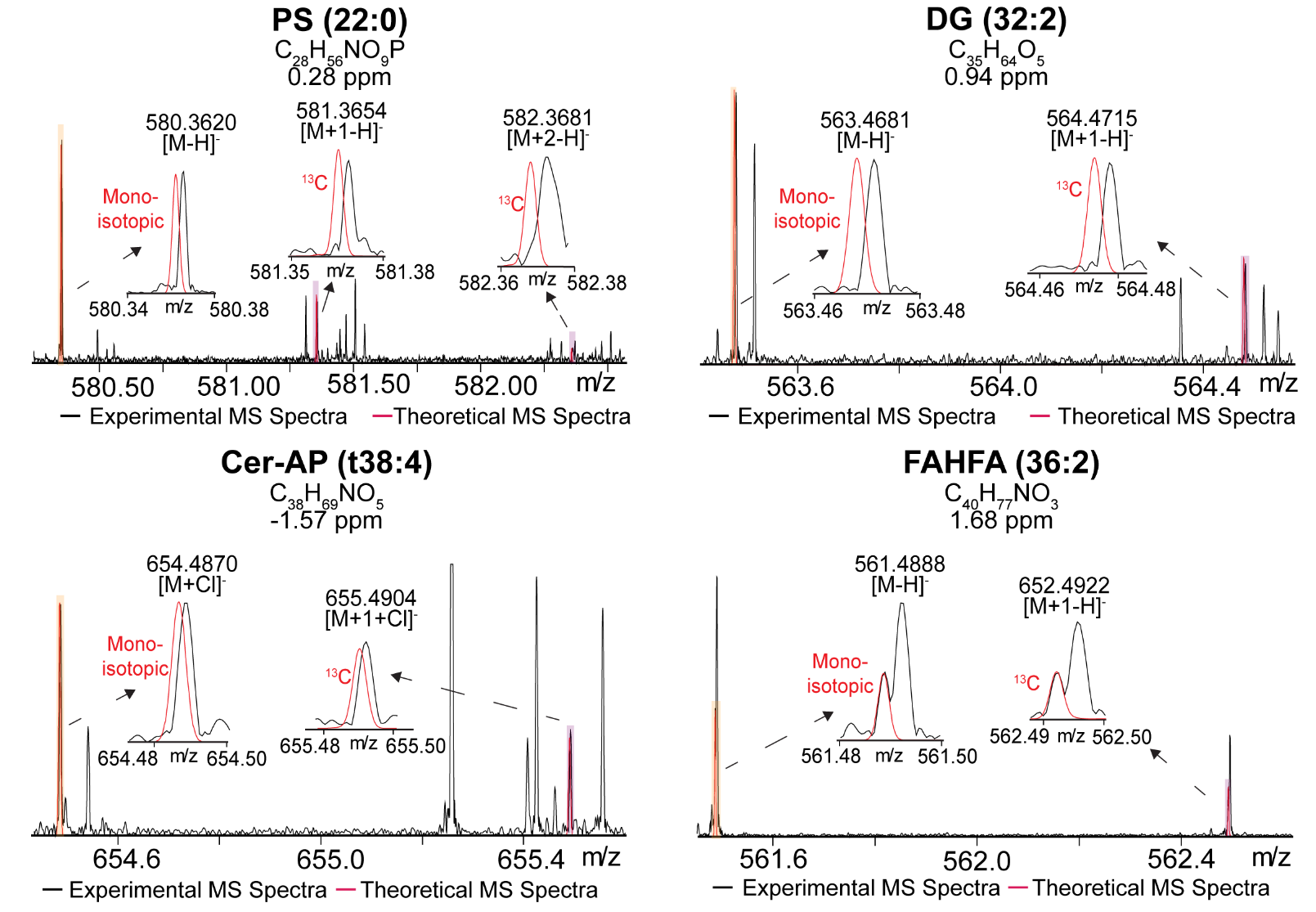

**Figure S8. Representative isotopic fine structure of newly reported metabolites in negative ion mode.** Shown are representative ultrahigh resolution DI-FTICR MS spectra in negative ion mode. Experimental mass spectra (black) were overlaid with theoretical mass spectra (red) to compare the isotopic distribution of newly reported metabolites. Cer-AP= ceramide alpha-hydroxy fatty acid-phytospingosine and FAHFA=fatty acyl ester of hydroxy fatty acid.

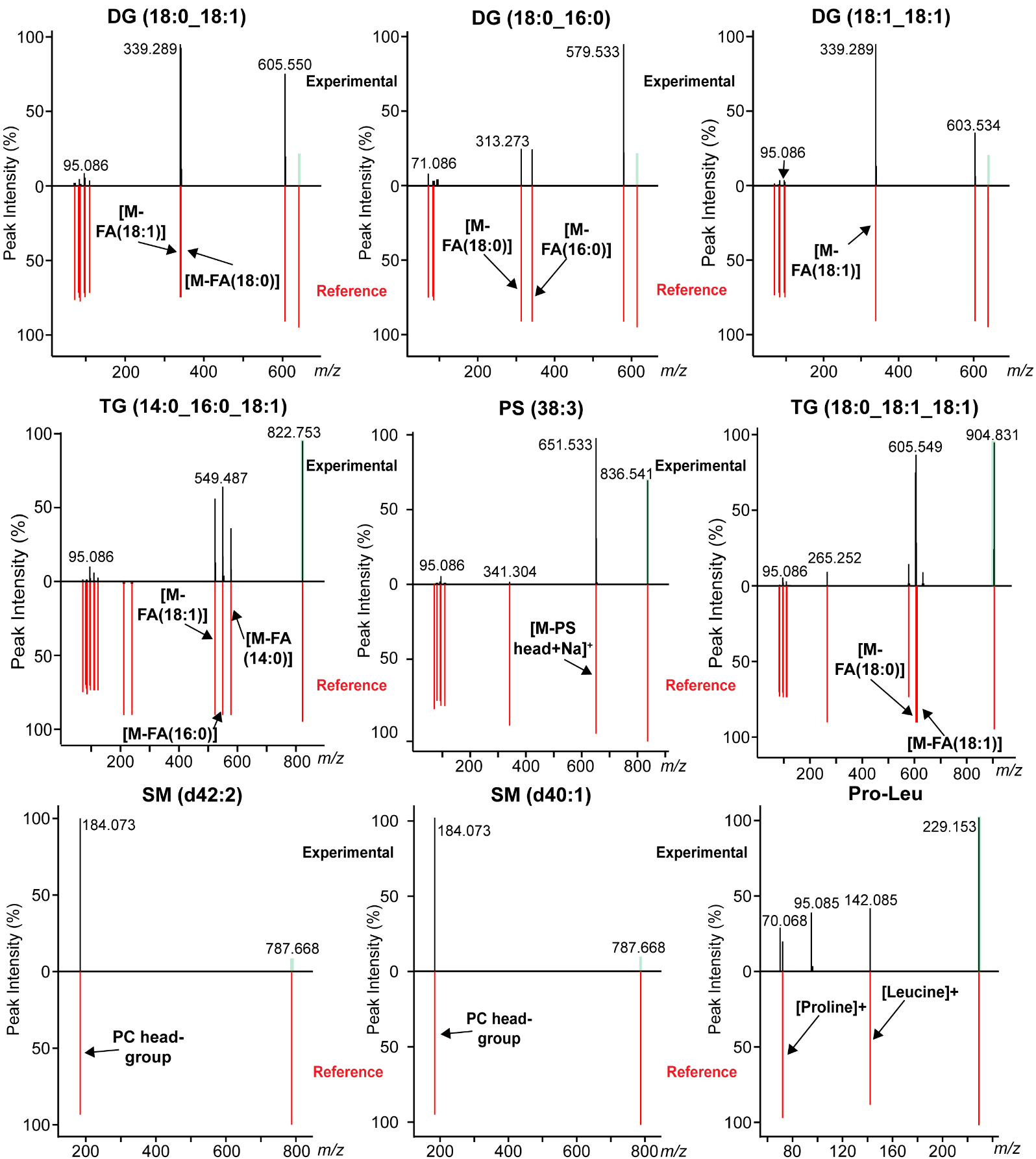

**Figure S9. Representative tandem mass spectra of newly reported metabolites in positive ion mode.** Representative positive ion mode tandem mass spectra from the LC-Q-TOF-MS/MS analysis. Shown are mirror plots comparing experimental mass spectra (black) with library (reference) mass spectra (red), which peak annotations. DG=diacylglyceride, TG=triacylglyceride, PS=phosphatidylserine. SM=sphongomyelin, and Pro-Leu=proline-leucine.

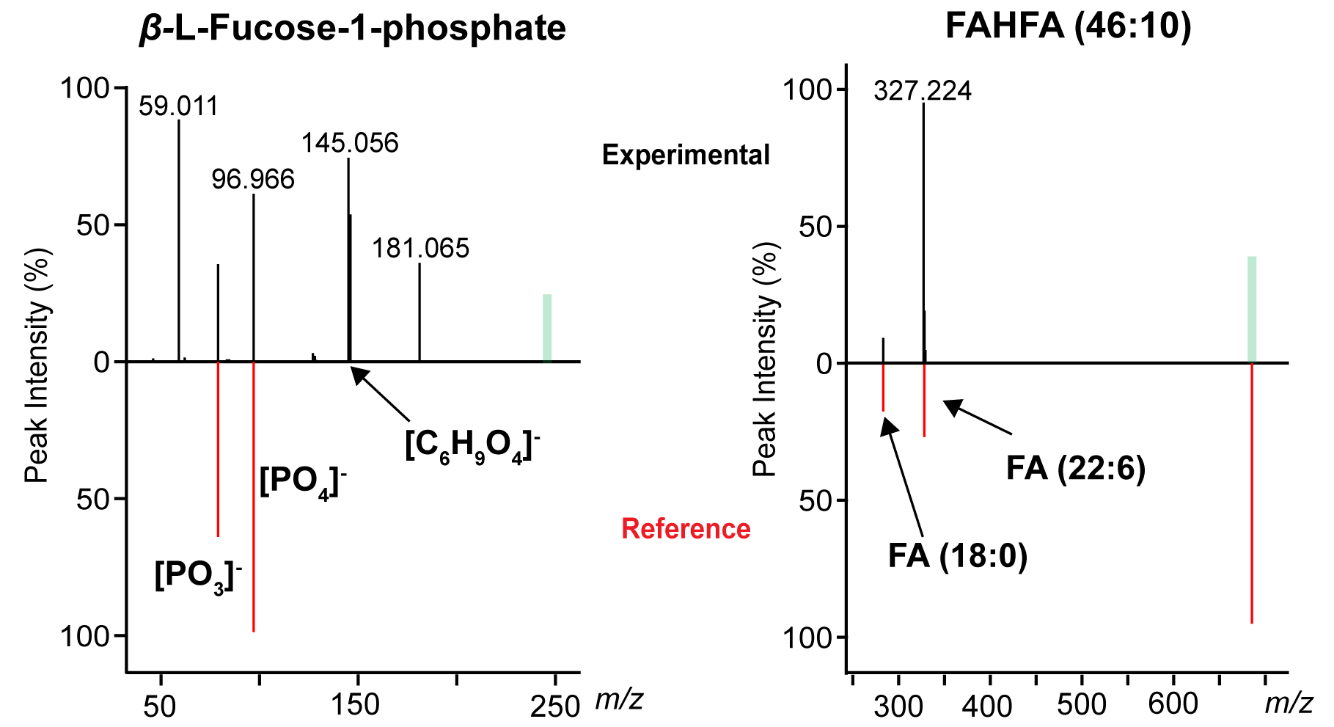

**Figure S10. Representative tandem mass spectra of newly reported metabolites in positive ion mode.** Representative negative ion mode tandem mass spectra from the LC-Q-TOF-MS/MS analysis. Shown are mirror plots comparing experimental mass spectra (black) with library (reference) mass spectra (red), which peak annotations. FAHFA=fatty acid hydroxy fatty acid.
